# A geometric-surface PDE model for cell-nucleus translocation through confinement

**DOI:** 10.64898/2025.12.18.695144

**Authors:** Francesca Ballatore, Anotida Madzvamuse, Cécile Jebane, Emmanuèle Helfer, Rachele Allena

## Abstract

Understanding how cells migrate through confined environments is crucial for elucidating fundamental biological processes, including cancer invasion, immune surveillance, and tissue morphogenesis. The nucleus, as the largest and stiffest cellular organelle, often limits cellular deformability, making it a key factor in migration through narrow pores or highly constrained spaces. In this work, we introduce a geometric surface partial differential equation (GS-PDE) model in which the cell plasma membrane and nuclear envelope are described as evolving energetic closed surfaces governed by force-balance equations. We replicate the results of a biophysical experiment, in which a microfluidic device is used to impose compressive stresses on cells by driving them through narrow microchannels under a controlled pressure gradient. The model is validated by reproducing cell entry into the microchannels. A parametric sensitivity analysis highlights the dominant influence of specific parameters, whose accurate estimation is essential to faithfully capture the experimental setup. We found that surface tension and confinement geometry emerge as key determinants of translocation efficiency. Although tailored to this specific setup for validation purposes, the framework is sufficiently general to be applied to a broad range of cell mechanics scenarios, providing a robust and flexible tool for investigating the interplay between cell mechanics and confinement. It also offers a solid foundation for future extensions integrating more complex biochemical processes such as active confined migration.

**Author summary:** Cells often migrate through very narrow spaces in tissues, a process critical for cancer invasion, immune surveillance, and tissue development. In particular, the stiffness of the nucleus, the largest and most rigid organelle, can limit migration through tight pores. In this study, we present a mathematical model describing the motion of a cell and its nucleus through a microchannel during cell translocation, using a geometric formulation based on surface partial differential equations. The model is general and applicable to a variety of scenarios involving confined cell transport. The model is validated by reproducing key experiments on cell translocation through narrow microchannels. The framework incorporates essential surface features, including mechanical responses, bending rigidity, and surface tension. Sensitivity analysis highlights surface tension and channel geometry as the parameters that most strongly influence translocation. Overall, the model provides new insights into the mechanics of confined cell transport, grants access to cellular quantities that are difficult to measure experimentally, such as cell and nucleus areas, perimeters, and stresses, and establishes a foundation for future extensions incorporating more complex biochemical processes.

## Introduction

Cell migration in confined environments is a fundamental mechanobiological process that plays a crucial role in many aspects of cellular life, including development and embryogenesis [1, 2], immune response [3, 4], and cancer metastasis [5, 6]. Cells may need to migrate through pores of varying sizes, which can sometimes be sub-cellular or even sub-nuclear. During migration through interstitial matrices, cells must overcome the physical barriers imposed by the dense extracellular network [7, 8]. To circumvent these constraints, cells secrete proteases that degrade the matrix, thereby enlarging pores and facilitating cell migration [9, 10]. Alternatively, cells that are unable to degrade the extracellular matrix (ECM) must squeeze through pre-existing pores in the matrix [11]. In this context, the role of the nucleus cannot be neglected, as it often represents the largest (ranging from 5 to 20 *µ*m) and stiffest organelle within the cell, acting as a physical barrier that can slow down movement or even inhibit migration altogether [12]. Cellular displacement under confinement is regulated by a complex interplay between biochemical signaling, cytoskeletal remodeling, and mechanical forces exerted by the ECM [13, 14]. Nevertheless, the precise mechanical response of cells to spatial confinement remains poorly understood. Recent studies have shown that cells adapt by modulating nuclear deformability, actomyosin contractility, and cytoskeletal organization to overcome geometric restrictions [15], indicating that confinement is not merely a passive barrier, but an active determinant of cell behavior and fate. Cells forced through microfluidic constrictions provide a valuable experimental platform for probing and quantifying their mechanical properties in vitro [16–19]. By observing how cells deform and cross narrow channels, researchers can gain insights into cellular stiffness, elasticity, and viscoelastic responses under controlled conditions.

To achieve a deeper understanding of this phenomenon, numerous mathematical models have been developed in recent years to quantify stress and strain fields and to identify the key parameters governing cellular responses to different stimuli [20]. Mathematical modeling serves as a powerful tool to investigate these processes, particularly when experimental data are scarce or difficult to obtain. Moreover, computational approaches are crucial for validating theoretical predictions against empirical observations, ultimately offering new perspectives on cell dynamics. The majority of these studies employ discrete approaches to examine cell invasiveness, focusing on parameters such as migration speed, displacement, and nuclear deformability. In [21], the authors investigate the role of nuclear deformability in regulating the ability of a cell to penetrate a three-dimensional extracellular structure, adopting a continuum mechanics description of the nucleus. Their findings emphasize the crucial interplay between the mechanical compliance of the nucleus and the cell’s capacity to form adhesive bonds, showing that both aspects jointly determine the efficiency of cell infiltration. In [22, 23], the authors employ a Cellular Potts Model to develop a computational framework for confined migration that explicitly accounts for both cell and nuclear sizes as well as their stiffness. These studies highlight the strength of multi-compartment modeling in unravelling the relative contributions of different physical factors that collectively shape migration under confinement. In particular, their results demonstrate that the size of the surrounding environment plays a decisive role in determining the observed migration phenotype. Mathematical models can also serve as powerful tools for designing microfluidic systems and organ-on-a-chip devices. For example, in [24], the authors establish a direct connection between computer-aided designs of microfluidic platforms and lattice-based frameworks for discrete cell modeling, enabling the simulation of collective cellular interactions. The entry of a cell into a microchannel has also been investigated in [25] using a Newtonian liquid-drop model, where the authors examine the translocation of breast cancer epithelial cells into a microchannel through both simulations and experiments. Their aim is to link the observed entry dynamics to the mechanical properties of the cells, thereby distinguishing between benign and malignant phenotypes. To model the interactions between cell deformability and fluid flow, coupled fluid–solid interaction frameworks have been widely adopted (see, for example, [26]). Through numerical simulations, these approaches describe how fluid forces (such as shear stresses and pressure gradients) deform cells and, conversely, how cell motion alters the surrounding flow. Typical numerical methods include, but are not limited to, finite element methods, immersed boundary methods [27], and Smoothed Particle Hydrodynamics (SPH) [28]. However, these formalisms predominantly focus on interactions between fluid flow and either rigid or weakly deformable moving boundaries, and often do not explicitly address the fully coupled mechanics of evolving cellular surfaces. On the other hand, [29] developed a hybrid agent-based/finite element model designed to capture distinct migration strategies. This multiscale framework for cancer cell motility integrates several key biological processes, including actin–polymerization-driven protrusions, cellular contractility, bleb formation, nuclear mechanics, and variable ECM attachment, as well as heterogeneity in ECM topology, thereby providing a comprehensive representation of cell migration dynamics under different conditions. Furthermore, coupled bulk–surface reaction–diffusion models for simulating single-cell migration have been developed in [30–32]. In particular, [31] demonstrated that the coupling between bulk and surface components allows a local perturbation on the cell membrane to generate propagating reaction waves, which eventually stabilize into a steady-state pattern due to the interaction with the bulk dynamics. In [32], a mechanobiochemical model for two-dimensional cell migration was proposed, coupling the mechanical properties of the cytosol with biochemical processes occurring near or on the plasma membrane. Moreover, [33] presented a conservative arbitrary Lagrangian–Eulerian (ALE) finite element method for the approximate solution of bulk–surface reaction–diffusion systems on evolving two-dimensional domains. Finally, the multiplicative decomposition of the deformation gradient was introduced in [34, 35] to account for both active and passive strains experienced by the cell. This mechanics-based computational approach successfully reproduces cell migration across different configurations, including migration under confinement through sub-nuclear microchannels, durotaxis, and locomotion on flat substrates. Taken together, these approaches have provided valuable insights into specific aspects of confined cell migration and translocation, yet their reliance on simplifying assumptions or their limited ability to simultaneously capture both cellular and nuclear mechanics leaves open important questions that motivate the development of complementary frameworks.

In this study, we adapt a geometric surface partial differential equation (GS-PDE) framework, previously developed in [36–39], to investigate cell translocation under confinement. Our choice of a GS-PDE framework is motivated by its ability to directly embed mechanical parameters within a geometrically consistent representation of both the cell plasma membrane and the nuclear envelope, thereby overcoming the limitations of previous models and enabling a more faithful reproduction of experimentally observed behaviours. The model is formulated in terms of evolution laws derived from a force balance principle applied to both the cell plasma membrane and the nuclear envelope. Within this framework, the cell plasma membrane and nuclear envelope are represented as dynamically evolving hypersurfaces, whose motion is governed by the interplay of multiple surface forces acting per unit area in the normal direction, as well as by their mutual interactions. The primary objective of this work is to assess how the mechanical properties of the cell and nucleus, together with the integrity of nucleus–cytoskeleton connections, influence cellular translocation in constrained environments. To achieve this, we aim to replicate the biophysical experiments described in [18], thereby validating the predictive capabilities of our mathematical model and reproducing key aspects of the observed cell behaviour. We aim to reproduce the physical and mechanical response of skin fibroblasts from healthy donors used as control cells in the experiments. Establishing this validation will enable the model to support further *in silico* investigations, providing deeper insight into the mechanistic basis of confined cell translocation and informing experimental design, thanks to the generality, applicability, and robustness of the modelling framework. Once validated in the context of confined transport, the model is then extended to predict the more complex process of confined active migration.

While we adopt the same general geometric and variational setting developed in [36–39], the focus and scope of the present study are substantially different. In particular, the novelty of this work lies in (i) the explicit coupling of two interacting evolving surfaces representing the cell plasma membrane and the nuclear envelope, (ii) the incorporation of experimentally calibrated mechanical parameters to reproduce confined cell translocation experiments, and (iii) a systematic sensitivity analysis identifying the dominant mechanical and geometrical factors governing translocation efficiency. Unlike previous GS-PDE models, which primarily considered single-surface dynamics or focused on abstract mechanical settings, the present framework is specifically designed to reproduce and interpret experimentally observed nuclear translocation under strong geometric confinement.

## Materials and methods

In this section, we first provide a brief overview of the experimental setup and measurements that we aim to model and later use for validation. Details of the experiments are described in [18]. Building upon this framework, we introduce the biophysical model and the governing equations developed to describe cell translocation through the microchannel. We then outline the numerical techniques employed to solve the model and discuss the key parameters that define the system’s behavior.

### Microfluidic experiments

We provide here a brief description of the biophysical experiments (Fig. 1), further details can be found in [18]. Microfluidic chips, fabricated from polydimethylsiloxane (PDMS), consist of a large quasi-two-dimensional flow channel (6-mm long, 0.3-mm wide, and 10 *µ*m-high to limits cells into one layer) with smaller microchannels in the middle (of 6 *×* 6 *µ*m^2^ squared cross-section and 100 *µ*m in length). Each chip held inlets to inject cells and buffer and an outlet that allowed to drive cells through the microchannels under controlled pressure gradient. Cells passing through microchannels were observed in brightfield microscopy and movies were acquired at frame rates varying from 125 to 1000 fps (see S1 Movie acquired at 500 fps as an example, corresponding to the cell timelapse shown in Fig. 1(b)). Experiments were carried out at 37°C, pressure gradient was fixed at 165 mbar. Movies were pre-processed using FIJI software to segment the cells, then their contour was analysed using a home-made Matlab routine to measure the temporal elongation of the cell tongue while they entered constrictions (see Fig. 1(c) corresponding to the cell shown in Fig. 1(b))).

**Fig 1.**
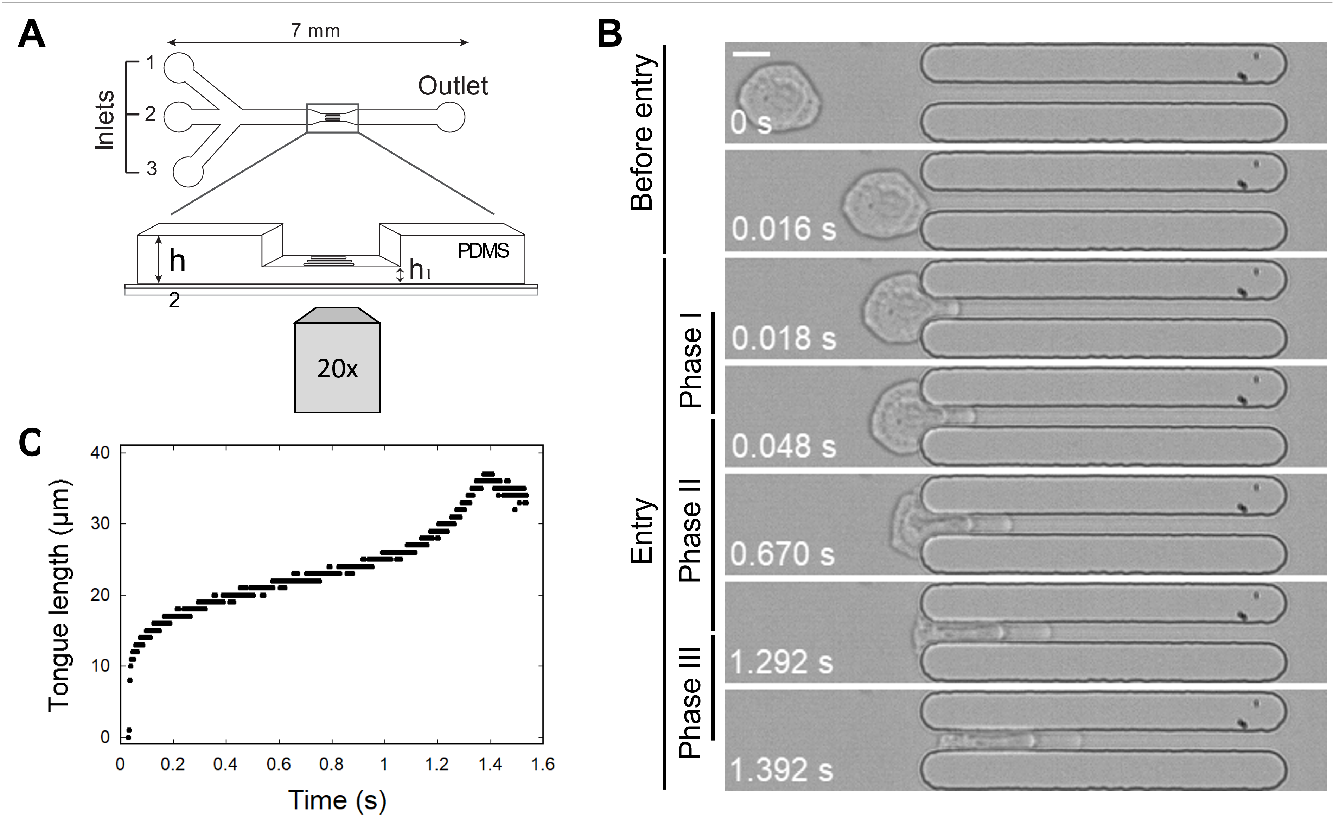
Biophysical experiment of cell translocation through microchannels. (a) Schematic top view of the microfluidic setup, with zoomed 3D-view of the microchannels. The cells passing through microchannels are observed from the bottom in brightfield microscopy. (b) Timelapse of a 15-µm cell passing through a microchannel. Time in sec. Scale bar: 10 µm. The timelapse is built from images extracted from S1 Movie. Entry phases I to III correspond to the three different dynamic regimes of cell tongue elongation shown in (c). (c) Cell tongue elongation in the microchannel over time, corresponding to the cell shown in (b). The cell was observed at a frame rate of 500 fps.

To quantitatively extract the cell mechanical properties from the microfluidic experiments, the measured tongue elongation was fitted using the Jeffreys rheological model. This model approximates the cell as a homogeneous viscoelastic material, characterizing its response through an effective elastic modulus, *E*, and two distinct short-time and long-time viscosities, *η*_1_ and *η*_2_. For cells used as reference for this work, the median values [and 95% confidence interval] of the mechanical parameters were *E* = 6 [5.6 − 6.3] kPa, *η*_1_ = 18 [17 − 21] Pa·s, and *η*_2_ = 277 [232 − 336] Pa·s. Further details on the mechanical parameters are given in Table 1.

**Table 1.** Reference parameters derived from biophysical experiments and literature.

| Par. | Value | Unit | Physical Meaning | Refs. |
| --- | --- | --- | --- | --- |
| $r_c$ | 13 | $\mu\text{m}$ | Cellular radius | [18] |
| $r_n$ | 5 | $\mu\text{m}$ | Nuclear radius | [18] |
| $w$ | 6 | $\mu\text{m}$ | Width of the micro-channel | [18] |
| $\tau$ | $10^{-3}$ | s | Time step of the simulation | estimated |
| $l_c$ | $5.0 \cdot 10^{-2}$ | $\mu\text{m}$ | Characteristic length | estimated |
| $k_s$ | 1.6 | nN/$\mu\text{m}$ | Surface tension | [48, 57, 58] |
| $k_b$ | $10^{-4}$ | nN$\cdot\mu\text{m}$ | Bending stiffness | [59] |
| $\omega$ | 0.1 | g/[($\mu\text{m}$)<sup>2</sup> · s] | Kinetic coefficient | estimated |
| $K_p$ | $1.2 \cdot 10^5$ | nN/($\mu\text{m}^2 \cdot \text{s}$) | Proportional gain | estimated |
| $K_i$ | 500 | 1/s | Integral gain | estimated |
| $D$ | 1.0 | kPa | Self-interaction strength | estimated |
| $\kappa_d$ | 5.0 | 1/$\mu\text{m}$ | Sharpness of the decay | estimated |
| $\delta_d$ | 0.1 | $\mu\text{m}$ | Minimal distance threshold | estimated |
| $\kappa_n$ | 5.0 | - | Normal orientation sensitivity | estimated |
| $\delta_n$ | -0.7 | - | Normal orientation threshold | estimated |
| $E_c$ | 0.0103 | kPa | Cell Young's modulus | [35, 61] |
| $E_n$ | 5.0 | kPa | Nucleus Young's modulus | [18] |
| $\eta_{1n}$ | 0.02 | kPa · s | Nucleus short-time viscosity | [18] |
| $\eta_{2n}$ | 0.3 | kPa · s | Nucleus long-time viscosity | [18] |
| $\eta_{1c}$ | 0.0002 | kPa · s | Cell short-time viscosity | estimated |
| $\eta_{2c}$ | 0.003 | kPa · s | Cell long-time viscosity | estimated |
| $\beta_1$ | 30 | kPa | Barrier force amplitude | estimated |
| $\beta_2$ | 3.5 | 1/$\mu\text{m}$ | Barrier force steepness | estimated |
| $F_{\text{pr}}$ | 16 | kPa | Pulling pressure acting on the cell | [18] |
| $k_{\text{rep}}$ | 12 | kPa | Cell-nucleus repulsive force amplitude | estimated |
| $\alpha$ | 5 | 1/$\mu\text{m}$ | Cell-nucleus repulsive force steepness | estimated |
| $k_{\text{el}}$ | 3 | kPa | Cell elastic restoring force amplitude | estimated |
| $k_n$ | 10 | kPa | Cell dragging force amplitude | estimated |
| $p$ | 4 | - | Cell dragging reduction factor | estimated |

### GS-PDEs for modeling the interplay between the cell plasma membrane and nuclear envelope during confined translocation

We consider a model for cell transport based on a geometric evolution law for two distinct yet coupled hypersurfaces, representing the cell plasma membrane and the nuclear envelope. Following the framework introduced in [37, 38], both surfaces evolve according to force balance equations per unit area, which incorporate the internal elastic response and the interaction forces between them. Specifically, we denote by Γ_*c*_ and Γ_*n*_ the evolving *C*^2^-smooth closed hypersurfaces, where the subscripts *c* and *n* refer to the cell plasma membrane and the nuclear envelope, respectively. Moreover, we consider here the case in which the cell passes through a microchannel under the action of an applied pulling pressure to reproduce the experimental conditions of [18]. A 2D projection of the initial configuration of the system is shown in Fig. 2. It must be noted that the mathematical formulation presented below is general and applies both to hypersurfaces embedded in three-dimensional space and to curves embedded in the two-dimensional plane.

**Fig 2.**
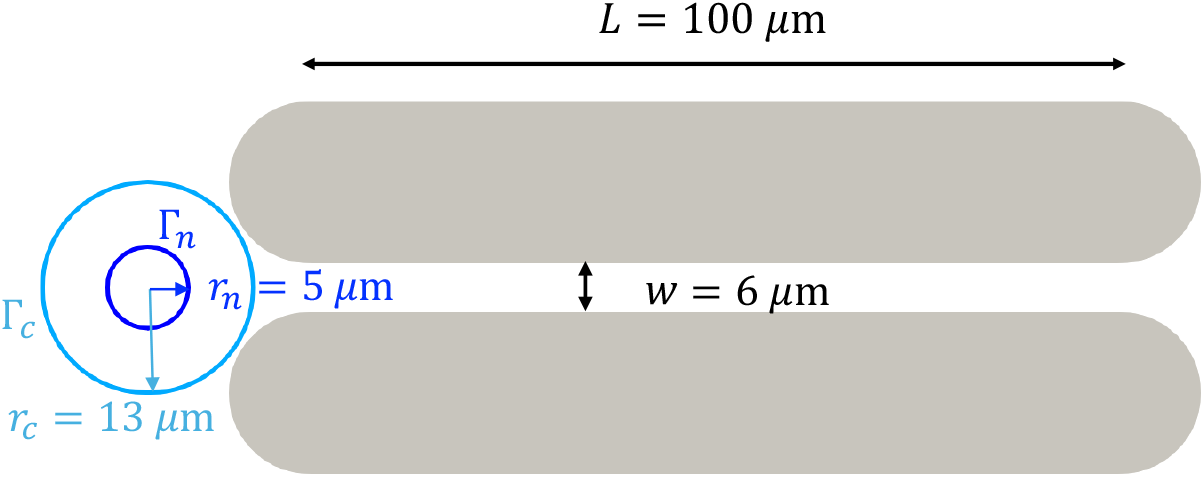
Schematic illustration of the initial configuration adopted in the 2D simulations to reproduce the 3D experimental conditions of [18]. The cell (light blue) and nucleus (dark blue) are placed at the entrance of a microchannel of width *w* = 6 *µ*m and length *L* = 100 *µ*m.

#### A geometric surface force balance equation for the cell plasma membrane

The force balance on the cell plasma membrane surface takes the form:

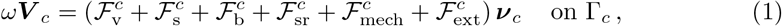

where ***ν***_*c*_ denotes the outward pointing unit normal to the surface Γ_*c*_ posed at each material point on the continuously deforming surface. Moreover, *ω >* 0 is a kinetic coefficient and ***V*** _*c*_ = *V*_*c*_ ***ν***_*c*_ denotes the material velocity of the cell plasma membrane. In the above, by definition 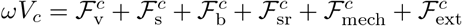. Since all force contributions are taken to act in the normal direction, the resulting membrane motion is purely normal by construction. Following established mathematical frameworks for geometric evolution equations (see e.g. [37, 40]), this modelling choice reflects two main assumptions: firstly, tangential velocity components correspond only to surface reparametrisations and do not affect the physical shape of the membrane; secondly, in the overdamped limit, any tangential reactive forces equilibrate instantaneously, allowing us to neglect explicit tangential tracking of the material points. The terms on the right-hand side of Equation (1) are described as follows.

- Experimental evidence indicates that, although the cell surface area can vary during translocation, the enclosed volume remains approximately constant [18, 41, 42]. We incorporate this observation as a hard constraint, implying that the cell can instantaneously compensate for small deviations in volume. We introduce a volume conservation model through the use of a penalty function, denoted by *λ*_*m*_(*t*), which can be interpreted as the time-dependent pressure difference between the cell interior and the surrounding environment. The resulting force associated with volume conservation is then expressed as

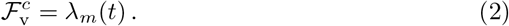

The associated force is written in vectorial form as *λ*_*m*_(*t*) ***ν***_*c*_, corresponding to the standard interface condition for a pressure acting on a deformable membrane. The parameter *λ*_*m*_(*t*) should not be interpreted as a strict Lagrange multiplier in the classical sense of constrained mechanics, even though in some studies it is referred to as a time-dependent Lagrange multiplier (see, for example, [32, 37]). Instead, it acts as a penalty pressure that softly enforces volume conservation through a relaxation mechanism. Its evolution introduces a controlled feedback that penalizes deviations from a reference volume, without enforcing the constraint exactly at all times. It is important to note that *λ*_*m*_(*t*) is spatially uniform and it satisfies the following first-order ordinary differential equation, corresponding to a proportional–integral control of the area:

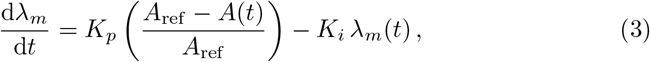

where *K*_*p*_ and *K*_*i*_ are positive constants representing the proportional and integral gains, respectively, and *A*(*t*) is the instantaneous cell area. The proportional term drives *λ*_*m*_ in response to the instantaneous deviation of the area from its reference value, while the integral term accumulates the effect of past deviations, ensuring long-term conservation. This formulation is similar to the approach originally proposed in [36, 43]. The parameters *K*_*p*_ and *K*_*i*_ are tuned to ensure stability and to achieve the desired level of accuracy in the area conservation. This penalty-based formulation allows for transient deviations from perfect volume conservation and reflects the fact that the constraint is enforced in an approximate rather than exact manner. From a biological perspective, this modelling choice is also justified by the ability of both the cytoplasm and the nucleus to transiently lose or acquire water during penetration into confined microchannels. As a result, cell and nuclear volumes are not expected to remain strictly constant throughout the translocation process.
- Resistance of the cell boundary to stretching may be incorporated by means of a surface energy of the form:

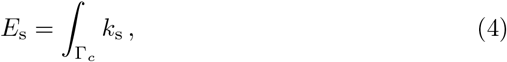

where *k*_s_ *>* 0 can be interpreted as a surface tension. The first variation of the surface area yields the mean curvature [44], hence the force arising from the surface energy is given by:

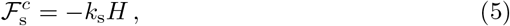

where *H* denotes the mean curvature of Γ_*c*_, understood here as the sum of the principal curvatures, i.e. 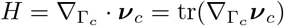. In the following, 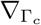 and 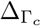 denote, respectively, the surface gradient and the Laplace–Beltrami operator, defined for an extension function *η* in a neighbourhood of Γ_*c*_ as

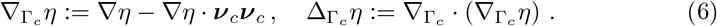
- The lipid bilayer forming the basic component of the cell plasma membrane also resists bending. We consider the established model of Helfrich [45] for the bending energy:

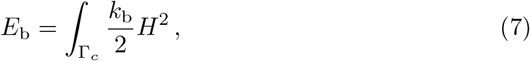

where *k*_b_ *>* 0 is the bending rigidity. The variation of the bending energy with respect to the surface yields the corresponding force contribution

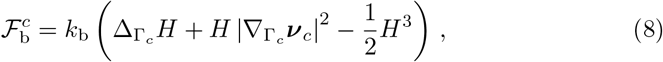

where | · | denotes the Frobenius norm, so that 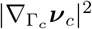 measures the local variation of the unit normal vector along the surface.
- To prevent non-physical self-intersections of the cell plasma membrane during large deformations, we introduce a self-repulsive force 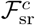 acting on each membrane point ***x***. This force is designed specifically to penalize configurations in which distant points on the membrane come excessively close and are oppositely oriented. In particular, for each membrane point ***x***, we evaluate its interaction with all other points ***y*** ≠ ***x*** that lie within a fixed distance threshold. The self-repulsive force at ***x*** is then defined as the maximum among these contributions:

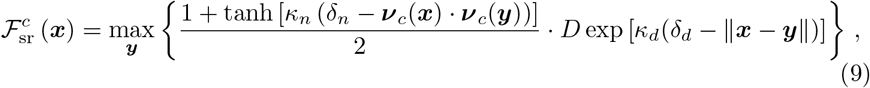

where ***ν***_*c*_(***x***) and ***ν***_*c*_(***y***) denote the unit outward normals at points ***x*** and ***y***, respectively, and ∥ · ∥ represents the Euclidean norm. The first factor reflects the alignment of the normals, becoming larger when the two normals are oppositely oriented. The second factor represents the proximity-based repulsion, which increases as the distance between the points decreases. The parameter *κ*_*n*_ controls the sensitivity to the relative orientation of the normals, and *δ*_*n*_ sets a threshold to detect near-opposing directions. The exponential term models the distance-dependent decay of the interaction, where *D* is the self-interaction strength per unit area, *κ*_*d*_ controls the sharpness of the decay and *δ*_*d*_ sets a threshold for the minimal distance between two points.
- We aim to capture and quantify the mechanical response of the cell surface within our model, enabling the direct integration of mechanical parameters estimated from biophysical experiments. Embedding mechanics within the evolving surface framework represents a significant advance, as it explicitly incorporates the mechanical response of the membrane-cortex composite. As the cell undergoes large deformations and translocation, a classical displacement-based elastic law is physically inconsistent due to the lack of a fixed reference configuration. Therefore, we model the cell surface as a deformable viscoelastic fluid (i.e. as an active viscoelastic surface), where the mechanical response arises from the velocity-dependent deformation of the surface. To ensure that deformations are restricted to the tangent plane, we define the surface rate-of-strain tensor 𝔻 using the surface projection operator ℙ = 𝕀 − ***ν***_*c*_ ⊗ ***ν***_*c*_ [46] as follows:

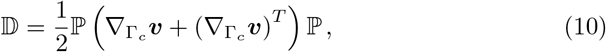

where ***v*** is the surface velocity. In our kinematic framework, where the velocity is assumed to act purely in the normal direction (***v*** = *V*_*c*_***ν***_*c*_), this expression simplifies to:

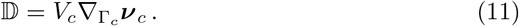

To describe the complex rheology of the actomyosin cortex, we adopt a surface viscoelastic constitutive law of the Jeffreys type. This model characterizes the cortex as a viscoelastic fluid, combining an instantaneous viscous response (denoted by ***σ***_*s*_) with a relaxing polymeric stress (denoted by ***σ***_*p*_). Hence, the total surface stress tensor ***σ*** is decomposed as ***σ*** = ***σ***_*s*_ + ***σ***_*p*_. The instantaneous viscous part ***σ***_*s*_ is given by:

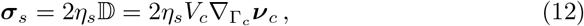

where 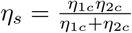 is the effective short-term cell plasma membrane viscosity, with *η*_1*c*_ and *η*_2*c*_ representing short- and long-term viscosities in the Jeffreys model. The polymeric part ***σ***_*p*_ represents the long-term viscoelastic relaxation and evolves according to:

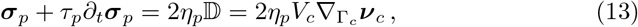

where 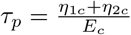 is the relaxation time, 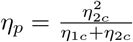 is the effective cell plasma membrane polymeric viscosity, and *E*_*c*_ is the surface Young’s modulus. A complete derivation of (13) can be found in Supporting material in Section S3. The mechanical force acting on the cell surface is computed as the normal component of the stress divergence. For a purely tangent and symmetric stress tensor ***σ***, this is rigorously given by [47]:

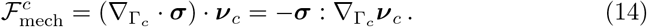

In our framework, the Helfrich bending term and the Jeffreys viscoelastic law are integrated to represent independent but mechanically complementary components of the cell boundary. The Helfrich term encodes the intrinsic resistance of the lipid bilayer to out-of-plane bending, acting as an instantaneous, purely geometric elastic constraint that penalizes highly curved shapes and prevents interface singularities. In parallel, the Jeffreys fluid formulation accounts for the active and dynamic in-plane rheology of the underlying actomyosin cortex. It decomposes the cortical response into two distinct physical timescales: an instantaneous viscous dissipation and a relaxing polymeric stress. The latter explicitly captures the transient elastic memory of the cross-linked actin network, which initially resists deformation but eventually yields and flows due to continuous filament turnover [48, 49].
- Finally, we consider the external forces 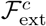 acting on the cell membrane, which, in our case, result from the sum of the following force contributions:

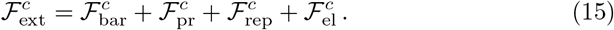
  – First, to model the interaction between the cell and external physical constraints, such as microchannel walls or obstacles, we introduce a barrier force 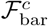 [37, 39]. This force prevents the cell from penetrating predefined regions by exerting a repulsive action whenever the membrane approaches specific barrier centres. We consider a cell passing through a microchannel of fixed width. The interaction with the barrier is modelled using an exponential repulsion force of the form

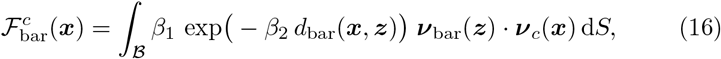

where ***z*** denotes points on the barrier surface, denoted by *B*. The coefficients *β*_1_ and *β*_2_ denote the amplitude and steepness of the cell plasma membrane and barrier interaction, respectively, and *d*_bar_(***x, z***) is the distance between the point ***x*** on the cell surface Γ_*c*_ and a point ***z*** on the barrier *B*. Finally, ***ν***_bar_(***z***) represents the outward-pointing normal vector at the barrier point ***z***.
  – Second, we introduce the force 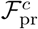 to account for the active pressure used to pull the cell in the microchannel. This force is modelled as a constant vector field applied in a fixed direction. In our model, we assume that this force acts uniformly in the positive *x*-direction, and it is defined as:

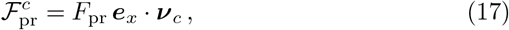

where *F*_pr_ is a constant magnitude and ***e***_*x*_ denotes the unit vector in the *x*-direction.
  – Moreover, we add the force 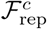, which ensures that the nucleus remains confined within the cell plasma membrane, specifically preventing it from crossing the cortical layer. This physical constraint is enforced via a penalization mechanism that introduces a repulsive interaction when the nucleus approaches the cortex. Let 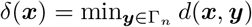 denote the minimal distance between a fixed point ***x*** ∈ Γ_*c*_ on the cell plasma membrane and all the points ***y*** ∈ Γ_*n*_ on the nuclear envelope. The repulsive force is defined as:

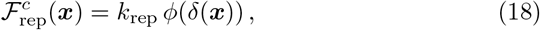

where *k*_rep_ *>* 0 is a stiffness parameter that determines the strength of the penalization and *ϕ*(*δ*) is a positive, increasing function as *δ* becomes smaller and closer to zero, for example:

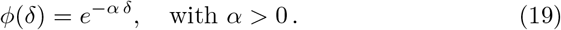
  – Lastly, we describe the interaction between the cell and the nucleus, due to the mechanical coupling via the cytoskeletal components, as an elastic force that prevents the cell plasma membrane from moving too far from the nucleus. For each point ***x*** on the cell plasma membrane, we compute the distance from the nuclear lamina, denoted by *δ*_el_. The resulting force included in our model is defined as:

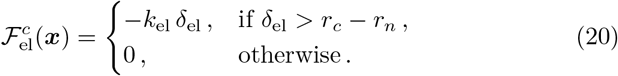

In the expression above, *k*_el_ is a constant parameter characterizing the stiffness of the interaction, *r*_*c*_ is the initial radius of the cell, and *r*_*n*_ is the initial radius of the nucleus. The mechanical coupling between the cell membrane and the nuclear envelope is modelled here through a single effective coupling parameter *k*_el_. We note that, at the biological level, force transmission between these two structures involves a complex network of cytoskeletal filaments, linker proteins (such as LINC complexes), and spatially heterogeneous attachments. The present choice is deliberately minimal and phenomenological. The parameter *k*_el_ should be interpreted as an effective mechanical coupling that aggregates the net contribution of these unresolved microscopic mechanisms at the cellular scale. This parsimonious formulation allows us to isolate and quantify the relative importance of membrane–nucleus coupling while keeping the model tractable and the number of free parameters limited. More detailed coupling mechanisms, including spatially heterogeneous, anisotropic, or non-linear interactions, could be incorporated within the same framework; we leave such extensions for future studies.

By combining all the contributions described above, the resulting balance of forces leads to the following governing 4th order GS-PDE for the evolution of the cell plasma membrane:

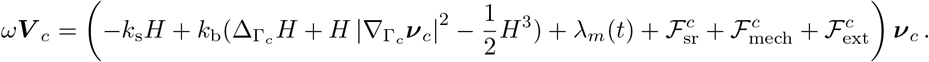

#### A geometric surface force balance equation of the nuclear envelope

The formulation is completed by introducing the surface force balance equation for the nuclear envelope. The nuclear envelope is subjected to its own set of forces, including contributions from volume conservation, viscous resistance, and mechanical stresses. In addition, external interactions with the cell plasma membrane are incorporated. The use of a surface-based mechanical formulation for the nuclear envelope deserves specific justification. While the governing equation for the nuclear envelope formally resembles that used for the cell plasma membrane, the underlying physical interpretation is different. The nuclear envelope is a composite structure consisting of a double lipid bilayer reinforced by the nuclear lamina, which is known to dominate the mechanical response of the nucleus at cellular length and time scales. Experimental studies have shown that, under confined migration, nuclear deformation is largely controlled by the mechanical properties of the lamina [50, 51]. As a result, modeling the nuclear envelope as an effective deformable surface provides a suitable description of the dominant mechanical resistance experienced during translocation through narrow constrictions. Within this effective framework, the surface tension, bending rigidity, and viscoelastic parameters associated with the nuclear envelope should be interpreted as phenomenological quantities that encode the integrated mechanical response of the lamina–membrane complex. Although the mathematical structure of the force balance is similar to that of the cell membrane, the corresponding parameters are chosen to reflect the significantly higher stiffness and resistance of the nucleus, consistent with experimental observations [52].

The overall balance of forces acting on the nuclear envelope in the normal direction is expressed as:

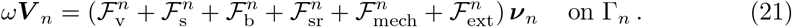

The main difference lies in the treatment of the external contributions, taking the following form:

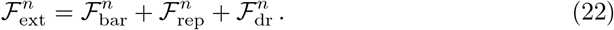

In this case, the nuclear envelope is subjected to the same barrier force, 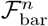, that also acts on the cell plasma membrane, as well as to a repulsive force 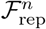 applied at every point of the nuclear surface. This repulsive force has the same functional form and parameters as the one acting on the cell plasma membrane, but with an opposite sign. In addition, we account for the dragging force, 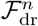. To ensure that the nucleus is carried along with the cell, we introduce a dragging force through which the plasma membrane transmits motion to the nucleus. This force represents the mechanical coupling between the cytoskeleton and the nucleus, ensuring that the nucleus follows the cell’s overall direction of movement. Its orientation is defined by the line connecting the cell center ***x***_*c*_ to the nuclear center ***x***_*n*_. Specifically, ***x***_*c*_ and ***x***_*n*_ correspond to the centers of mass of the cell and of the nucleus, respectively. Since only the normal components of the forces are considered in our framework, the interaction is projected onto the local normal vector of the nuclear surface. The resulting force is expressed as:

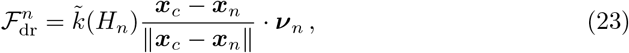

where 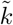 denotes the dragging coefficient, which depends on the curvature of the nucleus, *H*_*n*_. When the nucleus is undeformed or has fully traversed the microchannel, so that the cell is in a symmetrical configuration and the curvature at the front equals that at the rear, we set 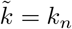. Conversely, during nuclear translocation through the microchannel, the ability of the cell to drag the nucleus is reduced, and we set 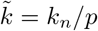, with *p >* 1.

### Numerical implementation

In the following, we present the variational formulation of the geometric surface evolution laws together with their corresponding finite element discretization. Since the models are almost identical except for some forces, we only present one variational formulation in a generalized form. Superscripts are omitted, and a unified framework valid for both cases is provided. We focus on the case of a curve embedded in the two-dimensional plane, corresponding to the biological problem we intend to model. The formulation can be naturally extended to three-dimensional evolving hypersurfaces. This extension is left for future studies.

The derivation is based on well-established results from differential geometry, reported in detail in [53]. We seek ***x*** ∈ *H*^1^(Γ(*t*))^2^ and *H* ∈ *H*^1^(Γ(*t*)), where *H*^1^(Γ(*t*)) denotes the Sobolev space of functions on the surface Γ(*t*) whose values and first derivatives are square-integrable and *t* represents the time, such that

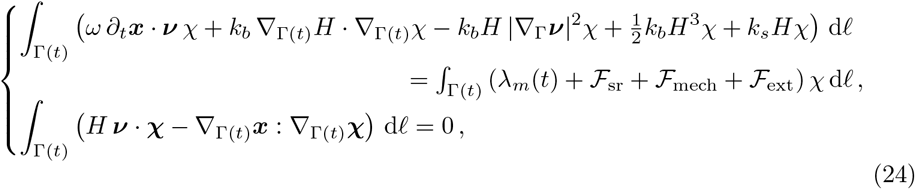

for all test functions *χ* ∈ *H*^1^(Γ(*t*)) and ***χ*** ∈ *H*^1^(Γ(*t*))^2^. Here, ***V*** = ∂_*t*_***x***, since we are working in a Lagrangian framework, where the material velocity coincides with the rate of change of the material points in time. In the above, we have called upon the Divergence Theorem and no boundary conditions since the surface has no boundary.

Then, a spatial and temporal discretization is performed, based on evolving surface finite elements [40, 54, 55]. We discretize the time interval [0, *T*] into *N* subintervals 0 *< t*_1_ *<* · · · *< t*_*N*_ = *T* with a fixed timestep *τ*. The method is based on parameterizing the surface 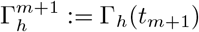 over 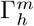. Given a discretised surface Γ_*h*_, we define the finite element space as

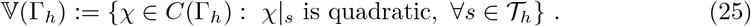

Then, for *n* = 0, …, *N* − 1, given 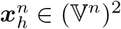 and 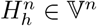, we seek 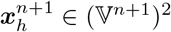 and 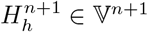 such that

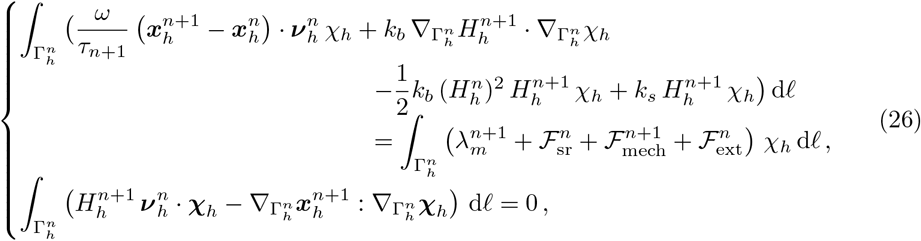

where “:” denotes the Frobenius inner product between two tensors. Here, we employ a Picard-type iteration to obtain a fully implicit backward Euler scheme.

To incorporate the surface rheology of both the actomyosin cortex and the nuclear envelope, the geometric evolution of each active interface is directly coupled with the 1D Jeffreys viscoelastic model. While the viscoelastic constitutive law is derived in a general tensorial form, our computational framework focuses on a 2D spatial setting where the cell and nuclear boundaries are modeled as 1D closed curves. In this lower-dimensional context, the surface tangent space is one-dimensional and any purely tangential tensor is fully characterized by a single scalar component *T*_*p*_. Specifically, the contraction of the shape operator and the polymeric stress tensor ***σ***_*p*_ reduces to the product *T*_*p*_*H*. For a complete derivation, we refer the reader to Section S3 of the Supporting Material.

Then, the normal mechanical force density ℱ_mech_ acting on either surface is evaluated implicitly at time *t*^*n*+1^ using the discrete curve variables:

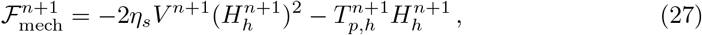

where 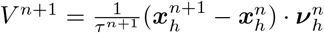 is the discrete normal velocity of the considered surface. Simultaneously, the scalar polymeric tension 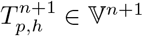 is updated dynamically by solving its corresponding weak formulation. Applying a fully implicit, unconditionally stable backward Euler time discretization to the viscoelastic evolution law, we seek to find 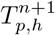 such that:

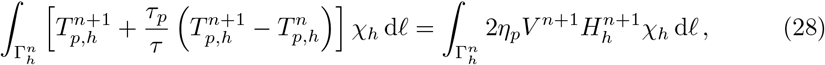

where *τ*_*p*_ represents the viscoelastic relaxation time of the specific component and *η*_*p*_ is its corresponding polymeric viscosity. This implicit sequential coupling ensures that the geometric stretching dynamically updates the active polymeric tension during the simulation, drastically improving numerical stability for both the cell and the nucleus.

Next, we proceed to compute the penalty parameter term *λ*_*m*_(*t*). Thus the penalty term 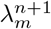, that enforces area conservation, is updated using the discrete version of proportional–integral scheme described in (3):

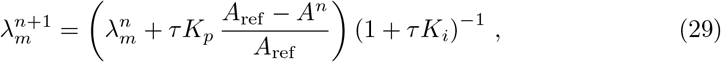

where *A*_ref_ denotes the reference initial area, *A*^*n*^ is the area at time step *n*, and *K*_*p*_ and *K*_*i*_ are the prescribed proportional and integral gains, respectively. It can be observed that, as the timestep approaches zero (*τ* → 0), the Lagrange multiplier remains effectively constant, i.e., 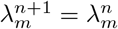.

From a numerical perspective, the minimal distance between the meshes of the plasma membrane and the nuclear envelope, required in Eqs. (18) and (20), is efficiently evaluated at each time step using a K-Dimensional Tree (KD-Tree) spatial indexing algorithm [56]. This approach drastically reduces the computational cost of the nearest-neighbor queries for the repulsive and attractive forces. Furthermore, the centers of the cell and the nucleus are computed as the geometric centroids of their respective meshes. Specifically, at each time step, the coordinates of the centers used in Eq. (23) are obtained by calculating the arithmetic mean of the spatial coordinates of all the corresponding mesh vertices.

### Parameter selection

Identifying and accurately estimating numerical parameters is essential for developing robust and reliable predictive models. However, many of these quantities are difficult to determine directly and must instead be inferred from in vitro measurements or estimated indirectly. In this work, parameter values were selected from previously published experimental and modelling studies, with particular reference to the experimental results reported by [18], so as to enable a direct comparison between our numerical simulations and their findings. The selected parameter values, together with the biological ranges found in the literature and the relative references, are summarized in Table 1.

First, we describe the geometrical dimensions of the cell and the microchannel, which are schematically illustrated in Fig. 2. Focusing on the size of the cell and its nucleus, we refer to the control skin fibroblasts reported in [18]. Accordingly, we consider an initial cell radius of *r*_*c*_ = 13 *µ*m and an initial nuclear radius of *r*_*n*_ = 5 *µ*m. As for the microchannel, we used the same width as in the reference paper [18], namely *w* = 6 *µ*m.

Second, in all numerical simulations, the following numerical parameters were adopted. The time step was set to *τ* = 10^−3^ s, while the characteristic length, corresponding to the typical size of the mesh elements, was chosen as *l*_*c*_ = 5.0 · 10^−2^ *µ*m for both the cell plasma membrane and the nuclear envelope. These values were selected to ensure numerical stability and convergence of the scheme, providing a suitable compromise between computational cost and accuracy.

Then, it is necessary to select appropriate values for the surface tension and the bending stiffness of the cell plasma membrane, since these parameters play a key role in determining cell shape and deformability under confinement. For the surface tension, we adopt the value *k*_*s*_ = 1.6 nN*/µ*m, which is consistent with the experimental estimates reported in [48, 57, 58]. For the bending stiffness, we use *k*_*b*_ = 10^−4^ nN · *µ*m, in agreement with the characterization provided in [59]. For the nuclear envelope, we increased both parameters by one order of magnitude to account for its higher rigidity.

Determining an appropriate value for the kinetic coefficient *ω* in the viscous force was not straightforward for either the cell or the nucleus, as only limited references are available in the literature. Following [60], and considering that this parameter directly influences the overall entry time of the cell into the microchannel, we calibrated it against the experimental data reported in [18]. On this basis, we set *ω* = 0.1 g/ (*µ*m)^2^ · s], a choice that can be generalized without loss of accuracy by a suitable rescaling of the time variable. Additional simulations performed with different values of *ω* confirmed its role in modulating the system dynamics. In the present study, we adopted the same value of *ω* for both the cell and the nucleus, although this assumption is not strictly necessary.

Regarding the proportional and integral gains *K*_*p*_ and *K*_*i*_ in Eq. (29), we tuned them to enforce a reasonable constraint on the change in cell area. Accordingly, we chose *K*_*p*_ = 1.2 · 10^5^ nN/(*µ*m^2^ · s) and *K*_*i*_ = 500 s^−1^.

For the self-repulsive force, all parameters were calibrated to reproduce the expected behavior. Specifically, we set the self-interaction strength per unit area to *A* = 1.0 kPa, the decay sharpness to *κ*_*d*_ = 5.0 *µ*m^−1^, and the threshold to *δ*_*d*_ = 0.1 *µ*m. In addition, the sensitivity to the relative orientation of the normals was set to *κ*_*n*_ = 5.0, with a corresponding threshold of *δ*_*n*_ = − 0.7.

We now consider the mechanical properties of both the cell plasma membrane and the nuclear envelope. For control fibroblasts, the Young’s modulus of the cell was estimated as a weighted average between a membrane stiffness of 100 Pa and a cytosolic stiffness of 10 Pa, yielding *E*_*c*_ = 0.0103 kPa. This choice is based on experimental measurements reported in [61] and on a reference value used in previous mathematical models [35]. For the nucleus, we adopted a reference elastic modulus of *E*_*n*_ = 5.0 kPa alongside nuclear envelope viscosities *η*_1*n*_ = 0.02 kPa · s and *η*_2*n*_ = 0.3 kPa · s, as estimated in [18]. Although these measurements were originally obtained for the cell as a whole, we attributed these properties directly to the nucleus, since the nucleus constitutes the stiffest and most mechanically resistant organelle. To account for the more fluid nature of the cell cortex, the corresponding cell plasma membrane viscosities were scaled down by two orders of magnitude relative to the values of the nuclear envelope viscosities.

Finally, we discuss the parameters associated with the external contributions. For the barrier force per unit area, we set the amplitude to *β*_1_ = 30 kPa and the steepness to *β*_2_ = 3.5 *µ*m^−1^. The pulling pressure acting on the cell was taken from the experiments reported in [18], with a value of 16 kPa. The penalization strength governing the repulsive interaction between the cell and the nucleus was set to *k*_rep_ = 12 kPa, with a steepness parameter *α* = 5 *µ*m^−1^. In addition, the elastic restoring force per unit area preventing the cell plasma membrane from moving too far from the nucleus was assigned a magnitude of *k*_el_ = 3 kPa, while the coupling strength enabling the cell to drag the nucleus during translocation was set to *k*_*n*_ = 10 kPa, with the parameter *p* fixed at 4.

## Results

In this section, we present the results of the proposed model, which successfully reproduce the biophysical experiments reported in [18]. In addition, we carry out a sensitivity analysis of key parameters involved in the model in order to assess how variations in their values affect the model predictions. This approach enables us to explore a broader range of scenarios and provides a means to evaluate the overall robustness and reliability of the model.

### Numerical model matches experimental results

In this section, we investigate the model’s ability to reproduce cell translocation within the microchannel, as well as the spatio-temporal evolution of the cell and nuclear shape deformations. This simulation employs the parameters listed in Table 1 and is designed to replicate the experiments reported in [18] and concisely summarized in the Microfluidic experiments section. Figure 3 illustrates the cell’s progression through the microchannel at various time points (see S2 Movie for the corresponding simulation). The simulation outcomes are in good agreement with the experimental observations presented in Fig. 1(b), where the passage of a cell through a microchannel is illustrated. To enable a more quantitative comparison, we examine the dynamics of cell entry into the constriction by analyzing the temporal evolution of the portion of the cell protruding inside the microchannel, hereafter referred to as the cell tongue like in [18]. The cell tongue length, evaluated at each time step, is reported in Fig. 4(a), showing a high similarity with the experimental curve from Fig. 1(c), replicated in Fig. 4(b) for facilitating the comparison. The comparison can be further processed looking closely on the dynamics of entry that can be divided into three successive phases. Phase I describes the cell entering the microchannel while the nucleus remains completely outside, Phase II corresponds to the nucleus entering the microchannel, and Phase III represents the cell passing through the microchannel once the nucleus is entirely inside. Each phase is characterized by a distinct quasi-linear increase in cell tongue length over time, i.e., a specific entry velocity. The computed velocities are *v*_I_ = 150.03 *µ*m/s, *v*_II_ = 17.23 *µ*m/s, and *v*_III_ = 111.25 *µ*m/s, corresponding to Phase I, II and III, respectively (cf. Fig. 4). The different regimes are in good agreement with the experimental findings reported in [18]: the same three phases were observed (see Fig. 4) and the measured values are of comparable order (see Table 2). It can be observed that in Phase II the nucleus slows down cell translocation, in agreement with experimental observations in other studies [62, 63]. Once both the cell and nucleus have fully entered the channel, the cell accelerates. This acceleration likely occurs because, once the nucleus has crossed the entry constriction, the cell experiences a more uniform confinement and reduced geometric resistance, allowing the driving forces to act more effectively and sustain a higher migration speed. It should be emphasized that the velocity values can vary substantially not only across different cell types, as expected, but also within the same cell type, as clearly reported by Jebane et al. [18] (see the confidence interval values in Table 2 derived from the dotplots shown in [18]). Such pronounced variability arises from the dispersion in cell size and mechanical parameters within a given population, as well as between different populations. For this reason, the slight discrepancies observed between our numerical and experimental velocities in Phases I and III, as shown in Table 2, fall within the experimentally observed variability and are therefore consistent with the reported data, as the values are of the same order of magnitude. As discussed in the Parameter selection section, the parameter *ω* can be tuned to reproduce different behaviours. For example, increasing *ω* by one order of magnitude relative to the previously adopted value leads to velocity reduction by one order of magnitude as well, namely *v*_I_ = 15.16 *µ*m/s, *v*_II_ = 2.26 *µ*m/s, and *v*_III_ = 11.45 *µ*m/s. It must be noted that the numerical simulation provides additional information on the position of the nucleus relative to the microchannel (see the vertical dotted lines in Fig. 4) which could not be measured in microfluidic experiments.

**Table 2.** Velocities of the three entry phases. Median and 95% confidence interval (CI) values were calculated from more than 400 cells across three independent experiment (see [18] for further details).

| Entry phase | Experimental velocity<br>Median [95% CI] ($\mu\text{m/s}$) | Numerical velocity<br>($\mu\text{m/s}$) |
| --- | --- | --- |
| Phase I | 483 [439–543] | 150.03 |
| Phase II | 20 [16–24] | 17.23 |
| Phase III | 214 [173–259] | 111.25 |

**Fig 3.**
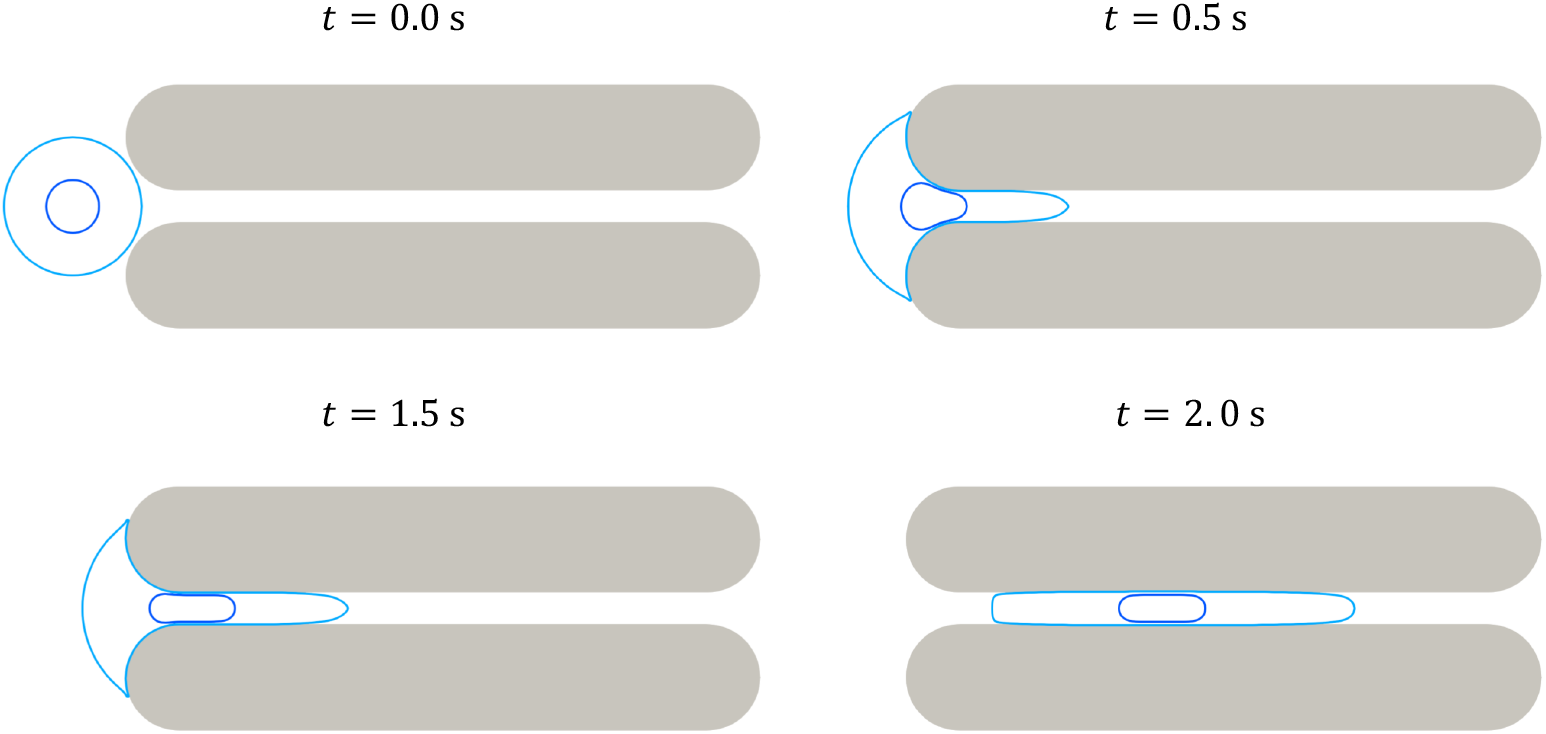
Snapshots of the cell plasma membrane (light blue closed curve) and the nuclear envelope (dark blue curve) at different time points for a microchannel width of 6 *µ*m. The simulation was carried out using the parameter values reported in Table 1.

**Fig 4.**
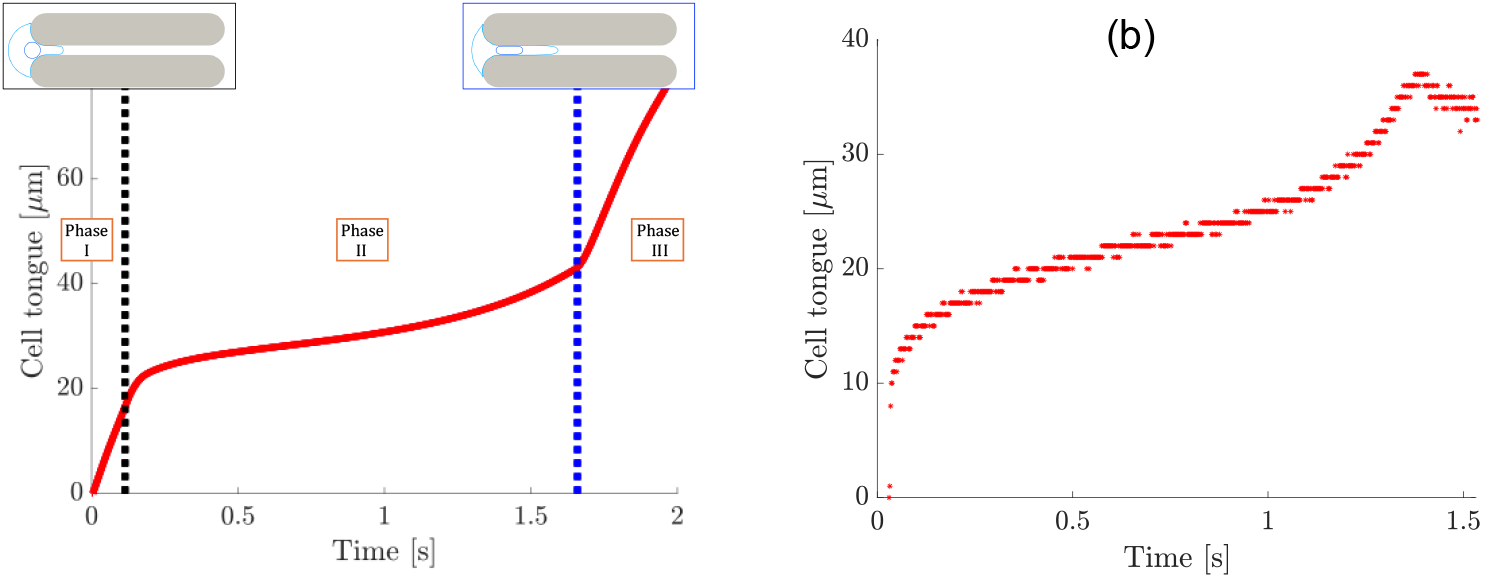
(a) Time evolution of the length of the cell tongue (red line). The black vertical dotted line marks the time at which the nucleus first contacts the microchannel, whereas the blue vertical dotted line indicates the time at which the nucleus is fully inside, as outlined in the snapshots above the lines. Phase I describes the cell entering the microchannel while the nucleus remains completely outside, Phase II corresponds to the nucleus entering the microchannel, and Phase III represents the cell passing through the microchannel once the nucleus is entirely inside. (b) Experimental curve reported from Fig. 1(c).

Figure 5 illustrates the temporal evolution of the principal quantities of interest. The figure highlights a key strength of our model: it provides access to quantities that are difficult to measure experimentally, including cell and nucleus area and perimeter, surface energy, and bending energy. These are computed directly from the evolving cell and nucleus surfaces in the simulation, offering valuable insights for experimentalists and enabling the investigation of biophysical properties that would otherwise be challenging to measure experimentally. In the first row, we show the time evolution of the total cell area (including the nucleus) and the nuclear area in Fig. 5(a), their perimeters in Fig. 5(b), and the velocity of the cell centroid during translocation in Fig. 5(c). All these quantities were computed from the discrete coordinates of the membrane and nuclear contours, by evaluating the enclosed area and the corresponding perimeter length. The area of the nucleus remains nearly constant throughout the simulation, whereas the area of the cell displays transient perturbations during entry into the microchannel and under full confinement due to compression. This indicates that the cytoplasm is softer than the nucleus, consistent with experimental observations [64, 65]. Quantitative measurements have shown that the nucleus is stiffer than the surrounding cytoplasm in many eukaryotic cells, with the latter allowing more pronounced area changes due to its lower stiffness. The cell perimeter exhibits a marked increase during the initial stage of the channel entry due to stretching and compression, consistent with the absence of perimeter constraints in the model. Variations in the perimeter of the nucleus are negligible, because the choice of the parameters for the nucleus makes it much stiffer than the cell plasma membrane. The centroid velocity shows a marked slowdown at the onset of cell penetration, followed by a further decrease as the nucleus enters the microchannel, since its higher rigidity hinders translocation. A partial recovery occurs once the cell has completely passed through the entry constriction, after which the velocity stabilizes. The second row shows the evolution of the surface energy and bending energy, defined in (4) and (7). They are shown in Fig. 5(d) and in Fig. 5(e), respectively. Regarding the surface energy, it exhibits a sharp increase at the beginning when the cell starts entering the microchannel, followed by a slower rise during nuclear translocation as the cell entry rate decreases, and finally a slight decrease once the cell is fully inside and reaches a steady shape. Similarly, the bending energy increases during the initial deformation of the cell as it adapts to the channel geometry, and then sharply decreases once the cell elongation becomes more uniform and curvature variations along the membrane are reduced.

**Fig 5.**
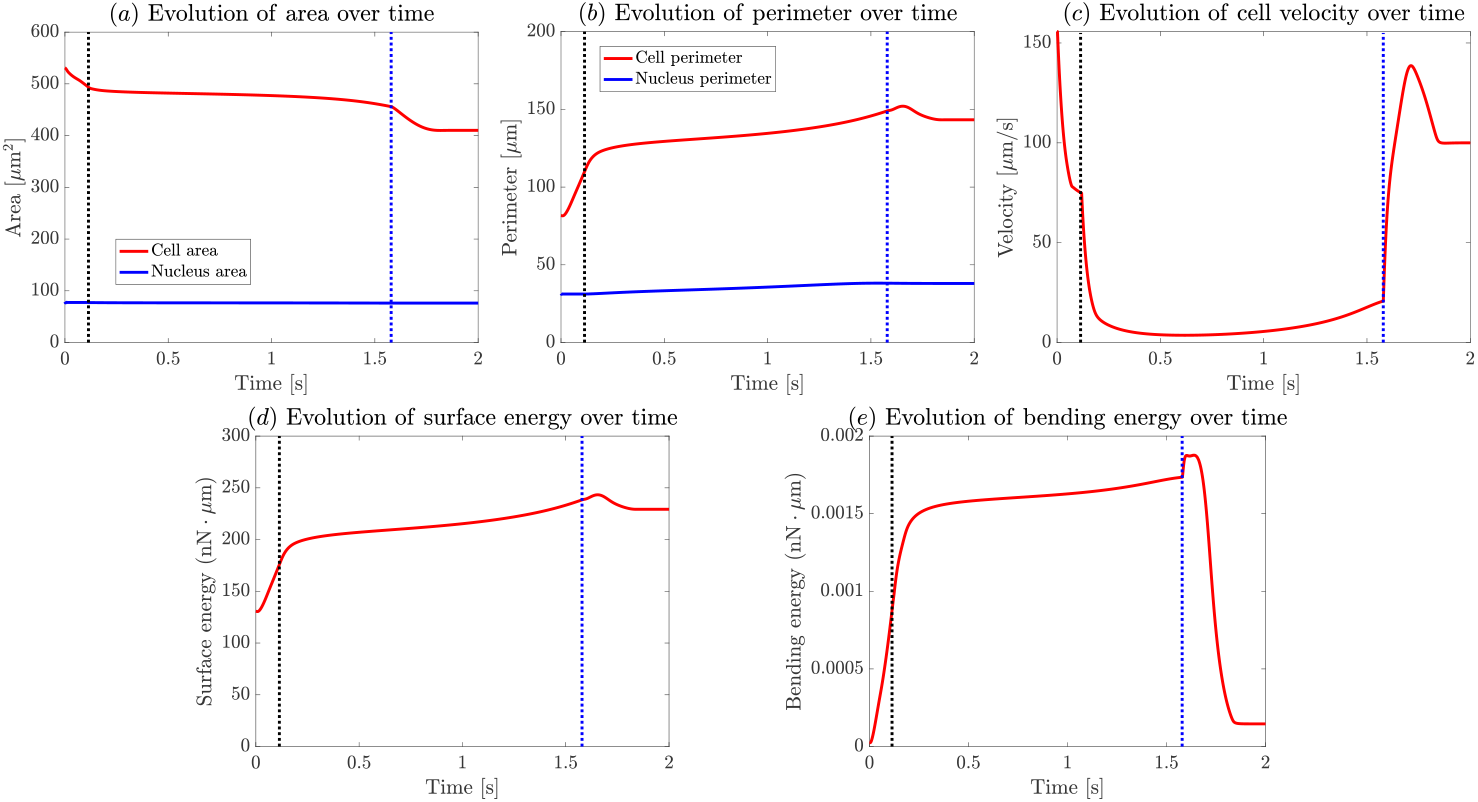
Time evolution of the key geometric quantities of interest. The first row shows the evolution of the cell and nucleus area and perimeter, as well as the velocity of the cell centroid. The second row shows the evolution of the cell’s surface energy and bending energy. The black dotted lines mark the time at which the nucleus first contacts the microchannel, and the blue dotted lines indicate when it is fully inside

We next reduce the microchannel width to *w* = 5 *µ*m in order to assess the behavior of the cell under a more severe geometric confinement. Using the mechanical parameters listed in Table 1, we find that the cell is unable to enter the channel and remains arrested at the entrance, as shown in Fig. 6. This response contrasts with the translocation observed in wider channels and indicates that the level of confinement has surpassed the cell’s ability to deform and squeeze through. In Fig. 7, the time evolution of the main geometric quantities of interest is reported. Regarding the area evolution in Fig. 7(a), a slight decrease in the cell area is observed as the cell initially attempts to enter the microchannel, after which it becomes arrested. A similar trend is seen for the perimeter in Fig. 7(b), which initially increases when the cell protrudes into the channel but then remains constant once the motion stops. No significant changes are observed in the nuclear area or perimeter. Concerning the velocity of the cell centroid in Fig. 7(c), a rapid decrease is observed compared to the initial value, until it eventually reaches zero when the cell is completely blocked at the channel entrance and no longer moves. Finally, for the surface energy in Fig. 7(d) and the bending energy in Fig. 7(e), we observe an initial increase followed by a plateau, as the cell shape and position remain unchanged after being trapped at the microchannel entrance. The results emphasize the pivotal role of geometric constriction in determining the predicted dynamics, which emerges as one of the two key factors governing cell translocation. The other crucial factor is the mechanical response of the cell itself: an accurate representation of stiffness and cortical tension is essential to capture whether a cell can adapt to extreme confinement or instead becomes mechanically trapped. Indeed, maintaining the same microchannel width of *w* = 5 *µ*m but halving both the nuclear Young’s modulus and surface tension allows the cell to deform sufficiently to pass through the constriction. These findings indicate that the nucleus acts as the primary mechanical barrier to cell translocation through narrow microchannels, and that softening its mechanical properties facilitates successful passage.

**Fig 6.**
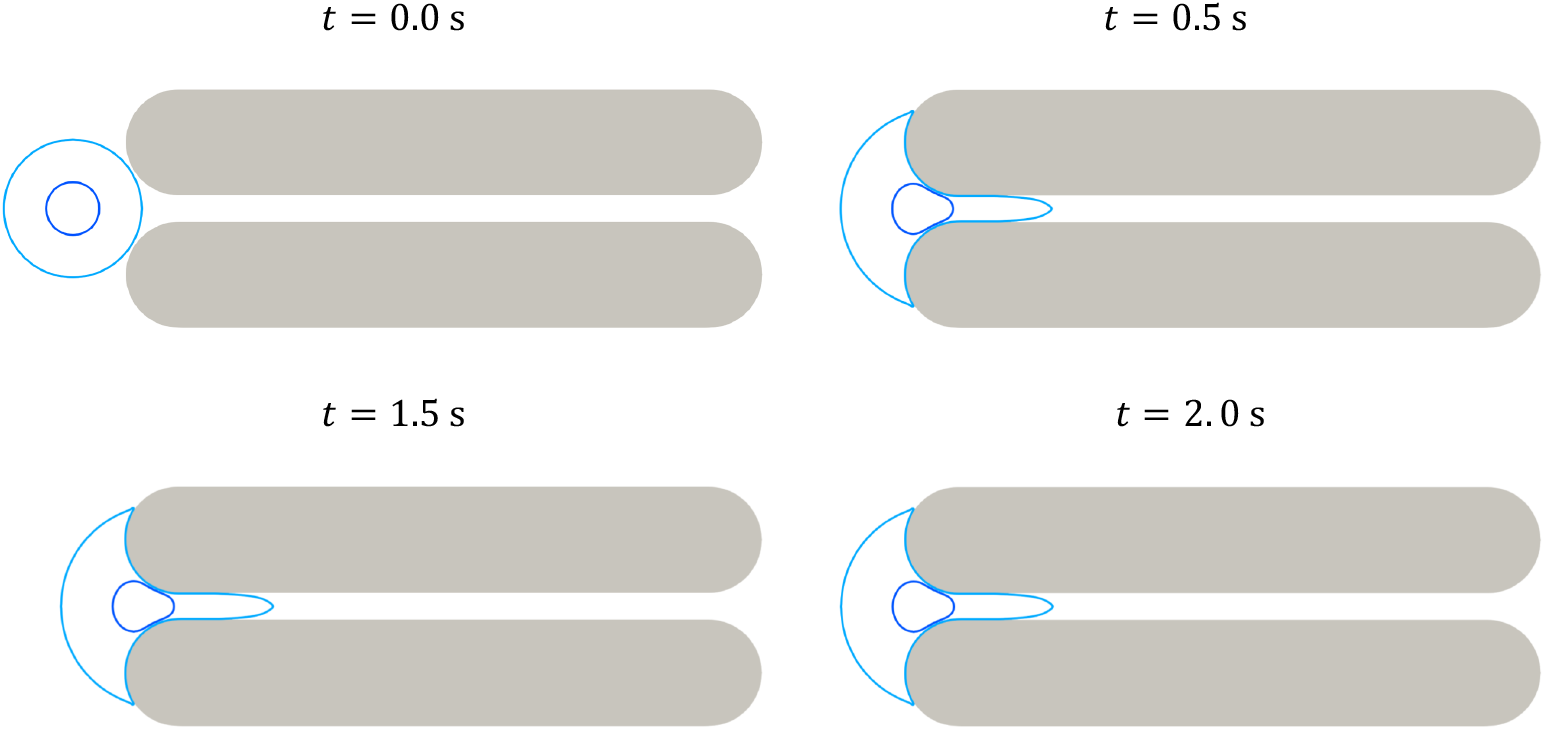
Snapshots of the space-time evolution of the cell plasma membrane (light blue closed curve) and the nuclear envelope (dark blue closed curve) at different time points for a microchannel width of *w* = 5 *µ*m. The simulation was carried out using the parameter values reported in Table 1 and a linear elastic constitutive law.

**Fig 7.**
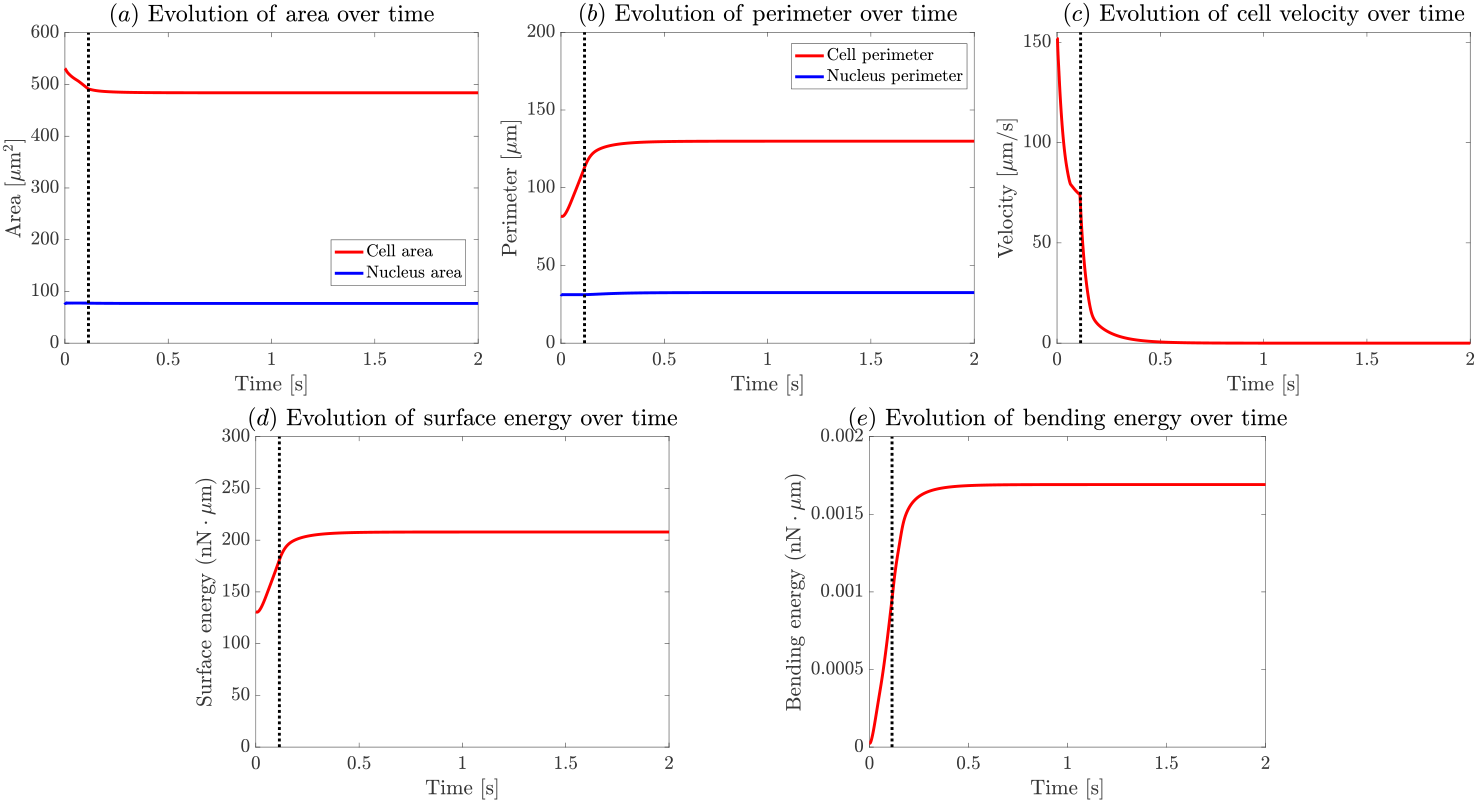
Time evolution of the main quantities of interest when the microchannel width is set to *w* = 5 *µ*m. The first row shows the evolution of the cell and nucleus area and perimeter, as well as the velocity of the cell centroid. The second row shows the evolution of the surface and the bending energy. The black dotted lines mark the time at which the nucleus first contacts the microchannel.

### Comparison between the viscoelastic and linear elastic models

In this section, we evaluate the model’s response by omitting the viscoelastic mechanical force defined in (14). The corresponding results are reported in Fig. 8. Compared to this purely elastic baseline, the full framework reveals that the primary effect of surface viscoelasticity is to attenuate the overall cell velocity, which consequently leads to an increased entry time. Fig. 9 illustrates the differences in cell centroid velocities and cell tongue length between the two cases during passage through the microchannel. Notably, the viscoelastic model exhibits a clear time-delay with respect to the purely elastic baseline: characteristic features such as the recovery of cell centroid velocity and the transitions in tongue length occur later in the viscoelastic case. Both graphs show the largest variation during Phase II, which depends strongly on nucleus size and mechanical parameters. The slower translocation observed in the viscoelastic case can be explained by the intrinsic dissipative nature of the material response. Unlike the purely elastic case, where the stress instantaneously follows deformation, viscoelasticity introduces time-dependent relaxation, causing part of the mechanical energy to be dissipated rather than stored. As a result, the cell deforms more slowly under the same driving forces, leading to a reduced velocity and a longer entry time. In particular, the viscous component opposes rapid shape changes, delaying both compression and recovery phases during transport through the constriction. Since this divergence is most pronounced when the nucleus begins entering the constriction, it indicates that a viscoelastic response of the nucleus has a greater impact on slowing translocation than applying viscoelasticity to the plasma membrane.

**Fig 8.**
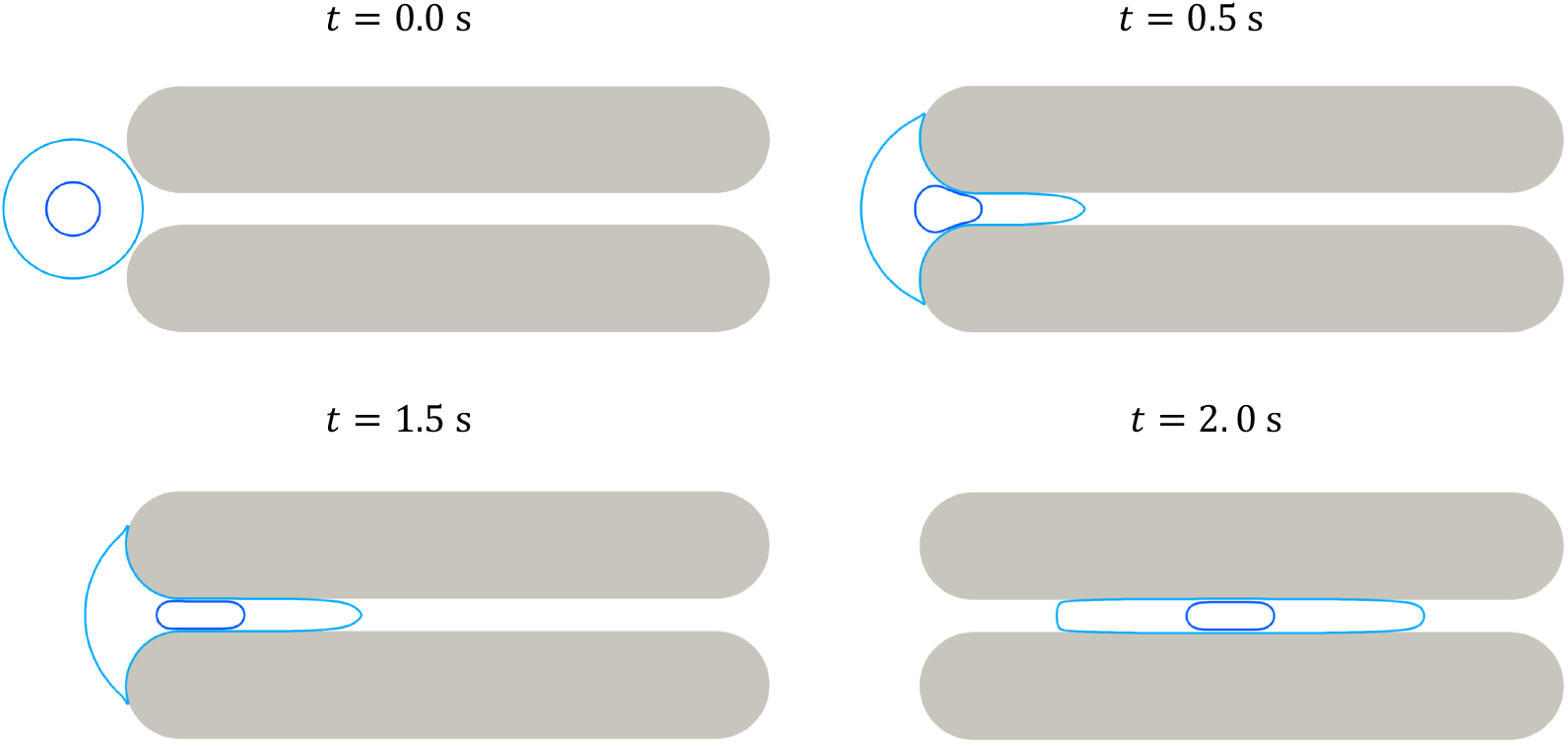
Snapshots of the space-time evolution of the cell plasma membrane (light blue closed curve) and the nuclear envelope (dark blue closed curve) at different time points for a microchannel width of 6 *µ*m. The simulation is performed using the parameters in Table 1, neglecting the viscoelastic mechanical force.

**Fig 9.**
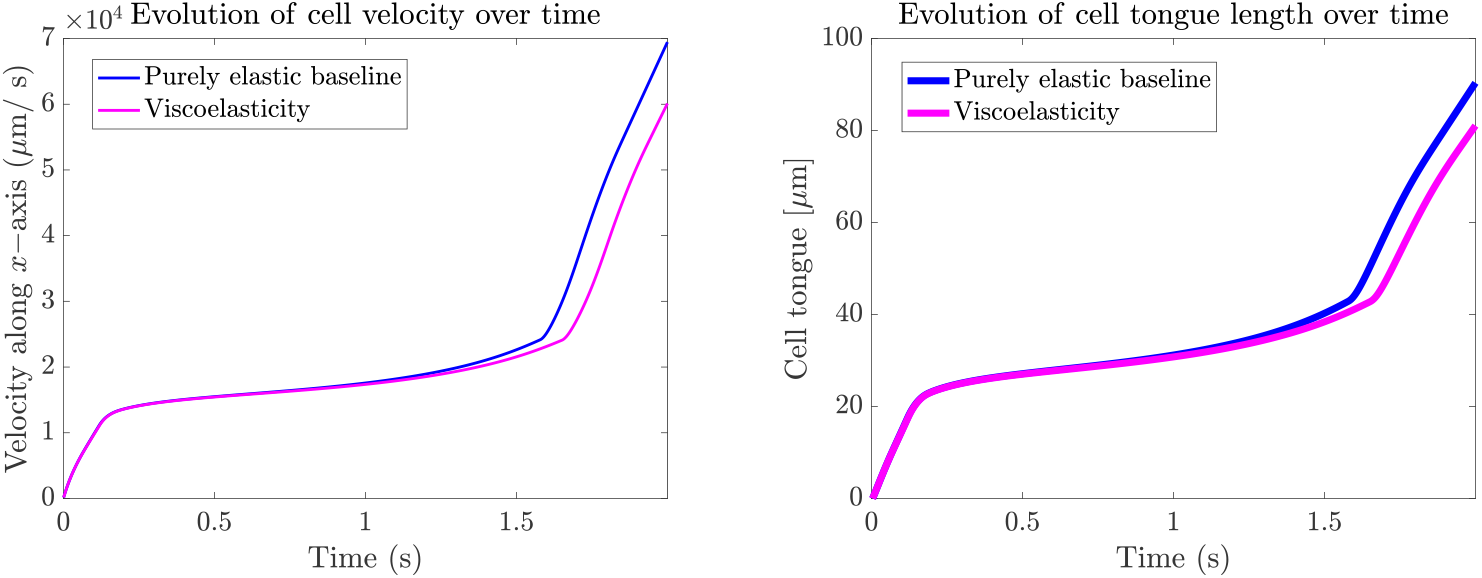
Comparison of the cell centroid velocity (left) and cell tongue length (right) between the purely elastic baseline and the viscoelastic case.

As a proof of concept, we verified that when the cell plasma membrane is assumed viscoelastic but not the nuclear envelope, the dynamics of the purely elastic baseline are essentially restored. Conversely, assuming a viscoelastic nucleus and a purely elastic plasma membrane recovers the full viscoelastic behaviour obtained in Fig. 3. Overall, the macroscopic effect of this structural dissipation is a reduction in cell velocity, leading to a quantifiable time-delay on the order of a fraction of a second in the total passage time.

### Sensitivity analysis

To assess the robustness of the proposed model and identify the parameters that most strongly influence cell dynamics under confinement, we performed a sensitivity analysis. The aim was to quantify how small perturbations in selected mechanical and geometrical parameters affect key observables associated with the translocation process. Specifically, we examined variations in the cell area, cell perimeter, and velocity of the cell centroid. Each parameter was independently perturbed by *±*10% around its reference value, as reported in Table 1, while all others were kept fixed. The parameters considered in this study include the cell Young’s modulus *E*_*c*_, the nucleus Young’s modulus *E*_*n*_, the microchannel width *w*, the pulling pressure acting on the cell *F*_pr_, and the surface tension *k*_*s*_ of both the cell and the nucleus.

Regarding the cell Young’s modulus, the corresponding results are presented in Fig. 10. Variations in this parameter exhibit a limited influence on both the cell and nucleus areas (see Fig. 10(a)), as well as on their perimeters (see Fig. 10(b)). Similarly, the cell velocity in Fig. 10(c) shows only minor changes. This is consistent with the experimental study where cells affected by a mutation of a component of the nuclear envelope, or by a biochemical treatment with a similar effect, did not display strong changes in the elastic modulus. This confirms that the Young’s modulus and the dynamics of confined transport are poorly related. Subsequently, we performed a similar set of simulations in which the cell Young’s modulus was fixed at its reference value, while the Young’s modulus of the nucleus, *E*_*n*_, was varied. No substantial differences were observed, and the results appear quite similar to those reported in Fig. 10.

**Fig 10.**
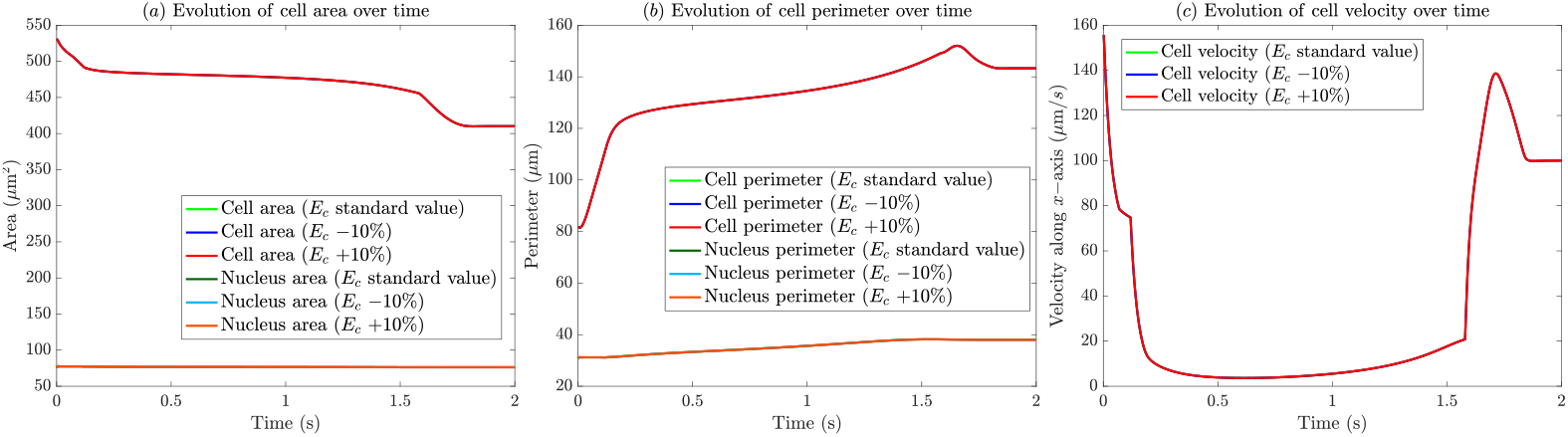
Sensitivity of (a) cell and nucleus areas, (b) perimeters, (c) centroid velocity, to variations in the cell Young’s modulus *E*_*c*_: standard value, −10%, and +10 %.

We next investigated the effect of a key geometrical parameter, the microchannel width *w*, which was previously identified as having a strong influence on cell passage. The results of this sensitivity analysis are shown in Fig. 11. Variations in *w* induce moderate changes in cell area (see Fig. 11(a)) and perimeter (see Fig. 11(b)); however, the most pronounced effect is observed on the cell velocity reported in Fig. 11(c). In particular, increasing the microchannel width enables the cell to traverse the constriction more rapidly, whereas reducing it causes the cell to remain trapped at the entrance within the simulated time frame.

**Fig 11.**
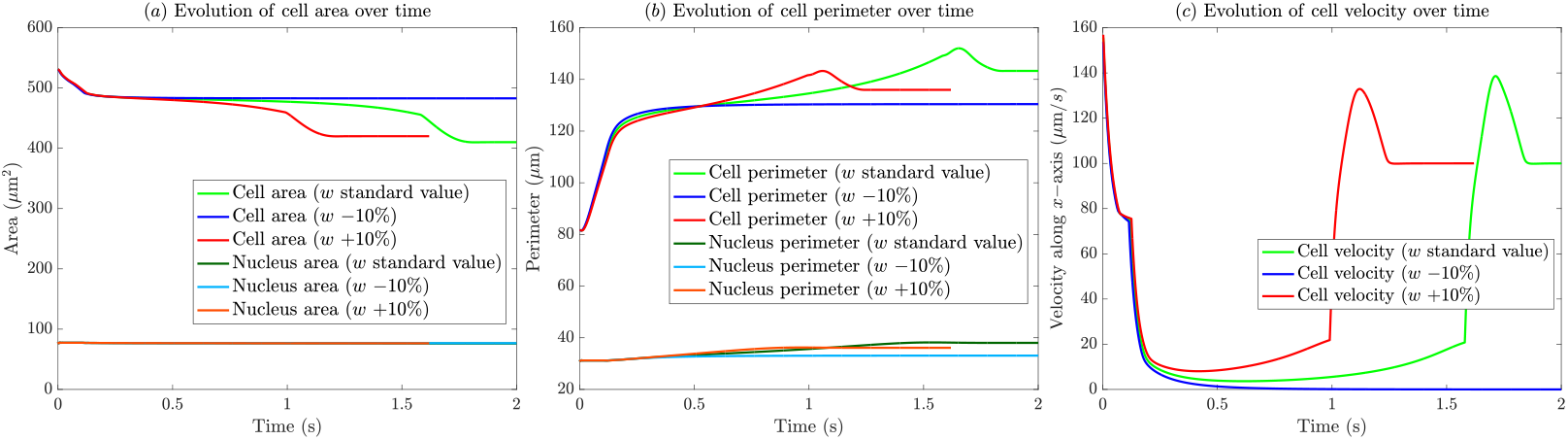
Sensitivity of (a) cell and nucleus areas, (b) perimeters, and (c) centroid velocity, to variations in the microchannel width *w*: standard value, −10%, and +10 %.

We also examined the influence of the pulling pressure acting on the cell *F*_pr_. The corresponding results are reported in Fig. 12. Overall, the variations induced by changes in this parameter are not substantial; however, slightly larger effects are observed in the cell perimeter in Fig. 12(b) and, more markedly, in the cell velocity in Fig. 12(c). In particular, after the nucleus has completely traversed the microchannel, a higher pulling pressure results in a higher instantaneous velocity of the cell.

**Fig 12.**
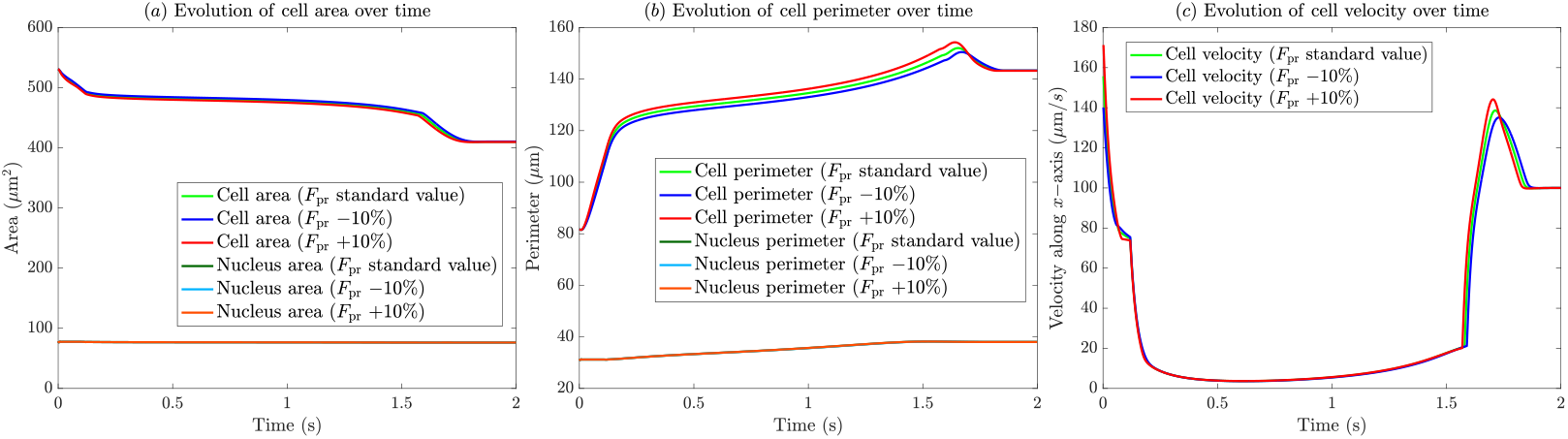
Sensitivity of (a) cell and nucleus areas, (b) perimeters, and (c) centroid velocity to variations in the pulling pressure acting on the cell *F*_pr_: standard value, −10%, and +10 %.

Finally, we considered the surface tension *k*_*s*_, which was varied simultaneously for both the cell and the nucleus. The results, presented in Fig. 13, reveal significant changes across all analysed quantities, confirming that surface tension is a key determinant of the model behaviour. Its influence is particularly evident for the nucleus, as it alters the time required for the nucleus to pass through the microchannel. This effect is especially visible in the velocity profile shown in Fig. 13(c), where the time at which the cell velocity increases, corresponding to the complete passage of the nucleus, depends on the value of *k*_*s*_.

**Fig 13.**
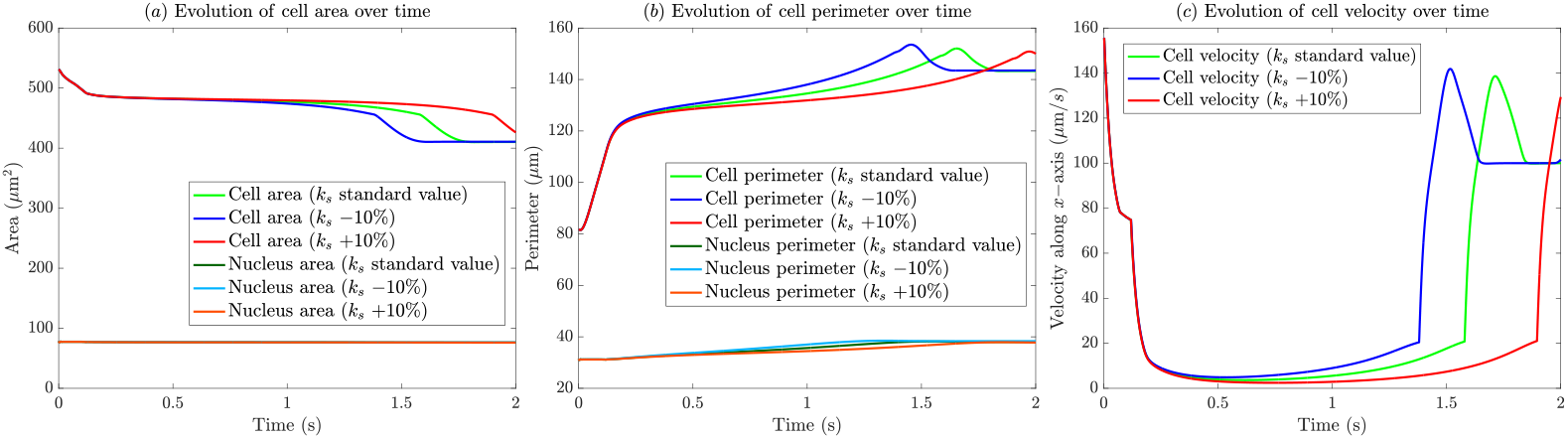
Sensitivity of (a) cell and nucleus areas, (b) perimeters, and (c) centroid velocity to variations in the surface tension *k*_*s*_: standard value, −10%, and +10 %.

Overall, this analysis highlights the parameters that most significantly affect the mechanical and transport response of the cell under confinement. In particular, surface tension and confinement geometry emerge as key determinants of transport efficiency, whereas cell stiffness primarily modulate local stress distributions. These findings provide valuable insight into the interplay between mechanical properties and geometric constraints, guiding future efforts in model calibration and experimental validation aimed at quantifying cellular and nuclear mechanics.

## Discussion and conclusion

In this work, we introduced a computational framework for simulating cell translocation under confinement. The proposed model successfully reproduces key experimental observations reported in [18]. Our main objective was to investigate the role of the nucleus, the stiffest organelle within the cell, during translocation through a narrow microchannel and to assess how its mechanical properties influence overall cell dynamics. The framework is based on geometric surface partial differential equations describing the evolution of both the plasma membrane and the nuclear envelope, as well as their mutual interactions and interactions with the surrounding environment. Each surface is characterized by specific mechanical properties, including surface tension, bending stiffness, Young’s modulus and viscosities. Model parameters were calibrated using experimental measurements from [18] and values available in the literature.

The model captures several key experimental trends, most notably the evolution of the cell tongue length, which exhibits three distinct phases. As the cell approaches the constriction, the tongue length initially increases, then slows when the nucleus reaches the microchannel entrance, and finally increases again once the nucleus fully enters the constriction. Overall, these results highlight that nuclear stiffness is a key regulator of translocation efficiency: a stiffer nucleus markedly hinders progression through confined environments, whereas a more compliant nucleus facilitates entry and passage. By contrast, cytoplasmic mechanics exert a comparatively secondary influence on the translocation process. Beyond reproducing measurable quantities, the model provides access to mechanical and geometric variables that are difficult to measure directly, such as the time evolution of the cell and nuclear area, perimeter, and the position of the nucleus relative to the microchannel. It also enables computation of surface energy, and bending energy, offering deeper insight into the mechanical state of the cell during translocation. Analysis of these quantities shows that, although total cell area is subject to a conservation constraint, it undergoes a slight reduction under compression, while the perimeter increases as the cell deforms to enter the constriction. Similar trends are observed for surface energy and bending energy, which sharply rise as the cell begins entering the microchannel, increase more slowly during nuclear translocation, and relax once the cell reaches a steady configuration within the channel.

A key aspect of the model concerns the mechanical characterization of the nucleus. In our study, we compared two constitutive formulations for the active interfaces: a purely elastic baseline and a surface viscoelastic model of the Jeffreys type. The comparison between these frameworks highlights the strong influence of surface rheology on the overall translocation dynamics. When the cell and the nucleus are modelled using the additional viscoelastic force, a dynamic dampening effect is introduced, which reduces the cell velocity and prolongs the translocation time. Consequently, both the cell and the nucleus resist rapid shape changes, delaying both the compression and recovery phases during passage through the constriction. The mechanical role of the nucleus is further evident when considering geometric confinement: reducing the microchannel width while keeping mechanical parameters constant causes the nucleus to become the limiting factor, blocking cell entry and preventing complete translocation.

Finally, a sensitivity analysis identified the parameters that most strongly influence translocation under confinement. Surface tension emerges as a major determinant of velocity, especially for the nucleus, directly affecting the time required for nuclear passage through the constriction. The degree of geometric confinement, namely the microchannel width, also strongly impacts translocation and can determine whether a cell can successfully pass or becomes mechanically trapped at entrance. Both parameters could in principle be systematically varied in the biophysical experiments, which would allow for a more comprehensive comparison between experimental observations and numerical simulations.

This study represents a first step toward a predictive mechanical model of single-cell translocation, developed within a surface-based spatiotemporal framework. The current setup serves as a validation benchmark, reproducing experimental observations and providing a robust foundation for future *in silico* studies. Future developments will incorporate biologically inspired cytosolic and membrane kinetics via surface reaction–diffusion equations, where state variables represent biomolecules regulating cellular protrusions, adhesion, and contractility. Inclusion of chemotactic cues will allow analysis of how cells sense and respond to spatial gradients under confinement.

Incorporating the extracellular matrix will allow us to explore translocation through heterogeneous, anisotropic environments, while extending the framework to multiple interacting cells will enable the study of collective transport and mechanical coupling between cells in confined geometries, also understanding tissue and organ formations. Finally, an important direction for future work concerns a more refined mechanical characterisation of the cell and nuclear membranes. In particular, recently developed models for viscoelastic surfaces could provide a more faithful description of cellular mechanical responses under large deformations and confinement, and therefore represent a natural extension of the present framework [46, 66–68].

## Acknowledgments

This work has been funded by the Agence National de la Recherche (ANR) through the MultiPhysC2M project (ANR-24-CE45-0690 to RA and EH) which also supports the postdoctoral fellowship of FB. The experimental work has been supported by Excellence Initiative of Aix-Marseille University - A*MIDEX (A-M-AAP-ID-17-66-170301-11.30 to EH): CJ performed and analyzed the microfluidic experiments under EH supervision. This work was supported by the INSMI Maths Vives International Research Program (awarded to RA and AM), Visiting Professor Campaign, Université Côte d’Azur (awarded to RA and AM), the Canada Research Chair (Tier 1) in Theoretical and Computational Biology (CRC-2022-00147 to AM), the Natural Sciences and Engineering Research Council of Canada (NSERC), Discovery Grants Program (RGPIN-2023-05231 to AM), the British Columbia Knowledge Development Fund (awarded to AM), Canada Foundation for Innovation – John R. Evans Leaders Fund – Partnerships (awarded to AM), and the British Columbia Foundation for Non Animal Research (awarded to AM). EH belongs to the French Consortium Approches Quantitatives du Vivant/Quantitative approaches to living systems (GDR AQV). FB is member of the National Group of Mathematical Physics (GNFM) of the Italian Institute for High Mathematics (INdAM). All funders had no role in the study design, data collection and analysis, decision to publish, or preparation of the manuscript.

## Supporting information

### S1 Movie

Movie of a 15-*µ*m cell passing through a 6 × 6 × 100 *µ*m^3^ microchannel, corresponding to Fig. 1(b-c). The images have been acquired at 500 fps.

### S2 Movie

Movie presenting the numerical simulation results of a cell passing through a microchannel of width 6 *µ*m, corresponding to Figs. 3-4.

### S3 The Jeffreys surface viscoelastic model

We model the cell surface using a tensorial Jeffreys fluid model, as done in [18] and for which we have the mechanical data. Rheologically, this is represented by a 1D circuit composed of two branches connected in series: a Kelvin-Voigt element (Branch 1) and a pure viscous dashpot (Branch 2), as described in Fig. SF1.

**Fig SF1.**
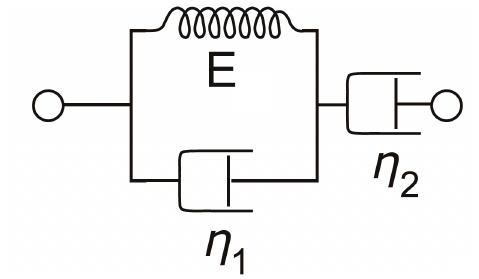
Jeffrey’s model described as 1D circuit composed of two branches connected in series: a Kelvin-Voigt element (Branch 1) and a pure viscous dashpot (Branch 2).

Because the elements are in series, the total surface stress tensor ***σ*** is uniform across both branches (***σ*** = ***σ***_1_ = ***σ***_2_), while the total surface rate-of-strain tensor 𝔻 is the sum of the individual strain rates (𝔻 = 𝔻_1_ + 𝔻_2_).

Branch 1 (the Kelvin-Voigt element) consists of a spring with elastic modulus *E* and a dashpot with viscosity *η*_1_ arranged in parallel. The stress in this branch is the sum of the elastic and viscous contributions:

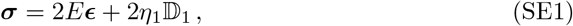

where ***ϵ*** is the surface strain tensor such that its objective time derivative yields the rate-of-strain (∂_*t*_***ϵ*** = 𝔻_1_).

In Branch 2 (the purely viscous dashpot), the stress is proportional to its rate-of-strain:

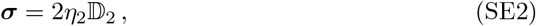

where *η*_2_ is the viscosity of the pure fluid component.

Taking the time derivative of the stress in Branch 1 (SE1) yields:

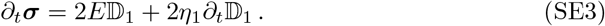

To obtain a single constitutive equation for the total stress, we substitute the kinematic compatibility condition 𝔻_1_ = 𝔻 - 𝔻_2_ into Equation (SE3):

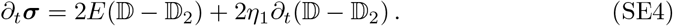

From the Branch 2 constitutive law (SE2), we know that 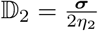. Substituting this into (SE4):

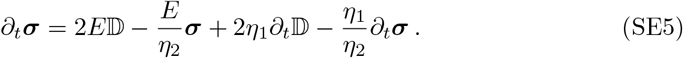

Rearranging the terms, we obtain:

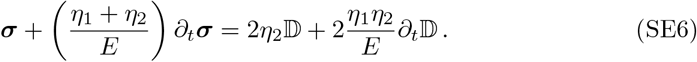

From a computational standpoint, computing the time derivative of the rate-of-strain (∂_*t*_𝔻) is numerically challenging because it requires tracking second-order time derivatives of the surface position. To elegantly bypass this, we decompose the total stress into an instantaneous viscous part (2*η*_*s*_𝔻) and a memory-dependent polymeric part (***σ***_*p*_) [69, 70]:

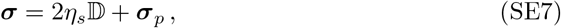

where we strategically define the effective instantaneous viscosity as 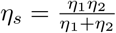.

Substituting this decomposition into Equation SE6 yields:

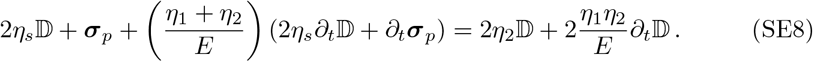

The terms involving ∂_*t*_𝔻 perfectly cancel out on both sides of the equation. This isolation provides a clean, and easily implementable evolution equation for the polymeric stress ***σ***_*p*_:

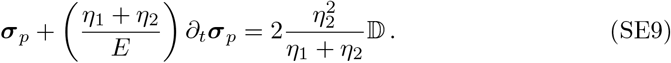

Then, we arrive at the final constitutive law governing the viscoelastic relaxation:

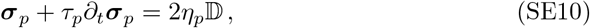

where 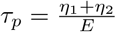 is the relaxation time and 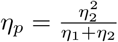 is the effective polymeric viscosity.

Finally, in the case one wants to model the cell as a 2D cross-section, the active surface reduces to a 1D closed contour. Consequently, the tensorial quantities defined on the manifold naturally simplify to their scalar counterparts along the curve. The polymeric stress tensor ***σ***_*p*_ reduces to the scalar macroscopic line tension *T*_*p*_, while the rate-of-strain tensor 𝔻 simplifies to *V H*, where *V* and *H* are the normal velocity and the curvature of the contour, respectively. The final constitutive law governing the viscoelastic relaxation thus becomes the scalar evolution equation:

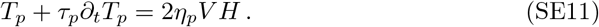

## Notes

https://github.com/francescaballatore/Constrained_cell_migration

### Competing Interest Statement

The authors have declared no competing interest.

### Summary of Updates

In this revision, we have carefully addressed some concerns raised and implemented several substantial changes.

